# Derepression of a single microRNA target causes female infertility in mice

**DOI:** 10.1101/2025.04.29.651088

**Authors:** Joanna Stefano, Lara E. Elcavage, Sue-Jean Hong, David P. Bartel, Benjamin Kleaveland

**Author notes:** These authors contributed equally to this work (Joanna Stefano, Lara E. Elcavage).

## Abstract

The miR-200a and miR-200b families control mouse ovulation and are essential for female fertility. The ZEB1 transcription factor is a conserved target of both families and has been implicated as a key player in female fertility at multiple levels. Using gene-edited mice that express a miR-200a/b-resistant form of *Zeb1*, we found that derepression of *Zeb1* in the female pituitary caused decreased production of luteinizing hormone and anovulatory infertility. These phenotypes were accompanied by widespread changes in pituitary gene expression characterized by decreased levels of ZEB1 targets, which include the miR-200a/b miRNAs, as expected from the miR-200a/b–ZEB1 double-negative feedback loop. Also observed were increased levels of mesenchymal genes, neuronal genes, and miR-200a/b targets. These results show that a double-negative feedback loop centered on the miRNA regulation of a single transcription factor can significantly influence the expression of thousands of genes and have dramatic phenotypic consequences.

## INTRODUCTION

Ovulation disorders are a common cause of female infertility (Carson and Kallen 2021; Munro et al. 2022). In mammals, ovulation relies on the coordinated synthesis and secretion of hormones from the hypothalamic-pituitary-ovarian axis. One of these hormones is luteinizing hormone (LH), a ∼25 kDa glycoprotein heterodimer comprised of a unique beta-subunit encoded by the *Lhb* gene and a common alpha-subunit that is shared with follicle-stimulating hormone (FSH), thyroid-stimulating hormone, and chorionic gonadotropin (Pierce and Parsons 1981). LH is secreted by gonadotrope cells of the anterior pituitary and stimulates follicle differentiation and steroid hormone biosynthesis in the ovary (Ma et al. 2004). During proestrus, an acute burst in LH secretion triggers release of one or more oocytes from the ovary and transforms the empty follicle into a corpus luteum, a structure that supports the fertilized oocyte until placentation occurs (Lunenfeld 2012; Szabó et al. 2024). Although mutations in *LHB* are a rare cause of hypogonadism and infertility in both women and men (Weiss et al. 1992; Valdes-Socin et al. 2004; Lofrano-Porto et al. 2007; Arnhold et al. 2009; Basciani et al. 2012; Yang et al. 2018; Song et al. 2018), abnormally low or high levels of LH are often associated with and likely contribute to female infertility arising from dysfunction of the hypothalamic-pituitary-ovarian axis (Mikhael et al. 2019). Given the central role of LH in female fertility and its dysregulation in fertility disorders, the regulatory processes governing LH expression and secretion are of substantial biomedical interest but remain incompletely understood.

A recurring motif in gene regulatory networks is the double-negative feedback loop (DNFL), which consists of two factors that inhibit each other (Alon 2007). By amplifying small changes to either factor, mutual inhibition generates a non-linear response to a linear stimulus, which allows some DNFLs to act as ultrasensitive “toggle” switches for bistable systems (Ferrell and Ha 2014). One of the best studied DNFLs centers on the miR-200a/b microRNA (miRNA) families and zinc finger E-box-binding homeobox 1 (ZEB1) and 2 (ZEB2) transcription factors (Korpal and Kang 2008; Bracken et al. 2024).

The evolutionarily related miR-200a and miR-200b families are post-transcriptional repressors that guide Argonaute proteins to the 3′ untranslated regions (UTRs) of target mRNAs, including *ZEB1* and *ZEB2,* which accelerates the deadenylation and decay of these mRNAs (Burk et al. 2008; Gregory et al. 2008; Korpal et al. 2008; Park et al. 2008; Bartel 2018). The ZEB1 and ZEB2 transcription factors bind CAGGTR consensus motifs in the promoters of hundreds of genes and inhibit or activate transcription, depending on the cellular context and interactions with co-repressors and co-activators (Heinz et al. 2010; Saitoh 2023). ZEB1/2 binding to the promoters of the *Mir200* loci inhibits transcription of the miR-200a/b host genes (Bracken et al. 2008; Burk et al. 2008), thereby establishing the miR-200a/b–ZEB1/2 DNFL.

The miR-200a/b–ZEB1/2 DNFL has wide-ranging functions but was first described in human cancer cell lines undergoing epithelial–mesenchymal transition (EMT) (Hurteau et al. 2007; Bracken et al. 2008; Burk et al. 2008; Gregory et al. 2008; Korpal et al. 2008; Park et al. 2008). In epithelial cells, a ZEB1^low^/miR-200a/b^high^ state maintains a polarized, stationary phenotype. This balance can be shifted by TGF-beta signaling and other factors that stimulate *Zeb1* transcription, leading to direct repression of key epithelial genes, such as *Mir200*, *Cdh1* (encoding E-cadherin), and *Epcam* (encoding epithelial cell adhesion molecule), and increased expression of mesenchymal markers such as vimentin and alpha-smooth muscle actin (Chua et al. 2007; Gregory et al. 2008; Korpal et al. 2008; Gregory et al. 2011; Kahlert et al. 2012; Zhang et al. 2016).

Besides its role in EMT, the miR-200a/b–ZEB1/2 DNFL has also been implicated as a key regulatory module in various aspects of reproductive physiology. During puberty, gonadotropin-releasing hormone (GnRH) neurons of the hypothalamus transition from a ZEB1^high^/miR-200a/b^low^ state to a ZEB1^low^/miR-200a/b^high^ state, which is thought to facilitate an increase in GnRH expression required for sexual maturation (Messina et al. 2016). An analogous transition has also been observed in the myometrial cells of the uterus during pregnancy; a ZEB1^high^/miR-200a/b^low^ quiescent state throughout gestation is converted to a ZEB1^low^/miR-200a/b^high^ contractile state at the end of pregnancy and during labor (Renthal et al. 2010). The ZEB1^low^/miR-200a/b^high^ state is also important in the pituitary, as female mice with either complete deletion of two miR-200b family miRNAs or forced *Zeb1* overexpression in gonadotropes are infertile due to reduced LH expression and a failure to ovulate (Hasuwa et al. 2013).

Although these experiments demonstrate critical roles for both miR-200a/b and ZEB1 in reproductive physiology, the particular importance of miR-200a/b-mediated repression of *Zeb1/2* or vice-versa is difficult to disentangle with traditional knockout or overexpression approaches because miR-200a/b and ZEB1/2 each have hundreds of targets. Here, we report on the molecular and physiological functions of the miR-200a/b–ZEB1/2 DNFL using *Zeb1* knock-in mice that are resistant to miR-200-mediated repression.

## RESULTS

### miR-200a/b represses *Zeb1* in the female pituitary

*Zeb1* and *Zeb2* are among the top predicted targets of the miR-200a/b families in both mice and humans (Fig. 1A,B; Supplemental Fig. S1A), owing to an unusually large number of conserved sites for each miRNA family residing in the 3′ UTRs of each gene (Agarwal et al. 2015). In addition, both *Zeb1* and *Zeb2* have been experimentally validated as miR-200a/b targets in diverse cancerous and noncancerous cells (Hurteau et al. 2007; Bracken et al. 2008; Burk et al. 2008; Gregory et al. 2008; Korpal et al. 2008; Park et al. 2008; Guan et al. 2018; Title et al. 2018). To investigate the physiologic role of miR-200a/b–mediated repression of *Zeb1* and *Zeb2* in mammals, we took advantage of the previously published *Zeb1* knock-in mouse (*Zeb1^200^*), which has point mutations in each of the nine predicted miR-200a/b binding sites in the *Zeb1* 3′ UTR (Fig. 1C), and also generated a *Zeb2* knock-in mouse (*Zeb2^200^*), replacing the wild-type *Zeb2* 3′ UTR with a mutant 3′ UTR in which either nine or ten of the 11 predicted miR-200a/b binding sites are each disrupted by 2–3 point mutations—thus creating a miR-200a/b–resistant *Zeb2* allele (Supplemental Fig. S1B,C). This knock-in approach offers advantages over more broadly used knockout and overexpression methods for interrogating the relationship between a miRNA and its target. Namely, this approach: 1) directly affects only one out of potentially hundreds of miRNA targets, 2) evaluates regulation of a target by all miRNA family members without requiring a priori knowledge about which miRNA family members are expressed in a particular cell/tissue, and 3) produces physiologically relevant increases in the target RNA and protein. The miR-200a/b–resistant *Zeb2* mice were viable and grossly normal (Table S1), and thus in this study, we focus on the phenotypic and molecular findings in the miR-200a/b–resistant *Zeb1* mice.

**Figure 1.**
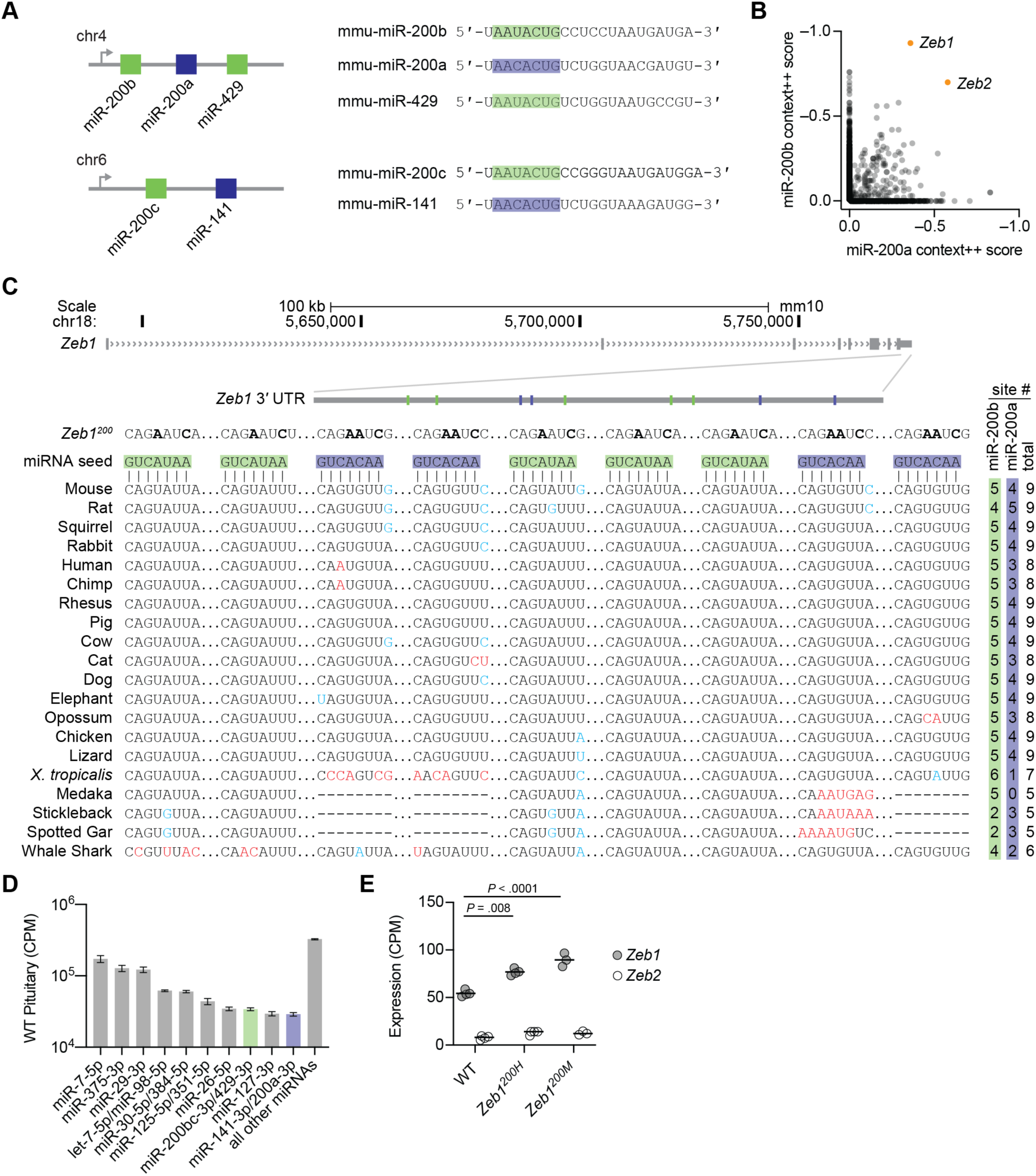
*Zeb1* is regulated by the miR-200 superfamily in the mouse pituitary. (A) The miR-200 superfamily members are expressed from two miRNA gene clusters. Diagrammed on the left are the miR-200b∼200a∼429 and miR-200c∼141 loci on murine chromosomes 4 and 6, respectively. Shown on the right are the mature sequences of each member of the miR-200 superfamily. The miR-200a/141 family seed sequence (blue) differs from the miR-200b/c/429 family seed sequence (green) by a single nucleotide at position 4 (counting from the 5ʹ end). (B) Predicted targets of miR-200b/c/429 and miR-200a/141 in mice. Plotted are Targetscan7 cumulative weighted context++ scores for both miRNA families with each circle representing a gene that is predicted as a target of miR-200b and/or miR-200a. Targets predicted with higher confidence have lower context++ scores. As expected for two miRNA families with distinct seed sequences, most predicted targets of miR-200b are not predicted targets of miR-200a and vice-versa. (C) Organization of the murine *Zeb1* locus. The *Zeb1* gene model (gray boxes, exons; > > >, introns) is depicted with an inset of the 3ʹ UTR indicating the location of the binding sites for miR-200b (green boxes) and miR-200a (blue boxes). Diagramed below that is the pairing of five miR-200b sites and four miR-200a sites and Multiz alignments for each site in 20 vertebrate species (red denotes a nucleotide change relative to the mouse sequence that disrupts pairing, blue denotes a nucleotide change that maintains at least a 7-mer seed site, some of which convert a miR-200b site to a miR-200a or vice-versa), as well the mutations (bold) that were introduced in each binding site to disrupt miRNA binding. The number of conserved miR-200b, miR-200a, and total sites are listed to the right of each alignment. (D) miRNA expression in wild-type pituitary. Plotted are the mean counts per million mapped miRNAs (CPM) of the ten highest expressed miRNAs and all remaining miRNAs, as determined by small-RNA sequencing, in wild-type (WT) pituitary from 28–36-week-old female mice (error bars, s.d.; n = 4). (E) Influence of *Zeb1^200^*alleles on *Zeb1* and *Zeb2* expression in the pituitary. Plotted are the counts per million mapped reads (CPM) for *Zeb1* and *Zeb2*, as determined by RNA sequencing, in WT, *Zeb1^200H^*, and *Zeb1^200M^*pituitary (black line, mean; n = 3–4 per genotype). Benjamini-Hochberg adjusted *p* values, as determined by a Wald test (DESeq2), are shown for *p* values less than 0.05.

To determine whether the mutant *Zeb1* allele was no longer susceptible to miR-200a/b-mediated repression, we examined *Zeb*1 expression in the female pituitary, a tissue with abundant miR-200a/b expression (Fig. 1D) (Hasuwa et al. 2013). RNA sequencing (RNA-seq) confirmed that disruption of miR-200a/b-mediated repression of *Zeb1* led to modest but significant increases in *Zeb1* mRNA—1.3-fold in heterozygous knock-in mice (*Zeb1^200H^*) and 1.5-fold in homozygous knock-in mice (*Zeb1^200M^*)—without affecting RNA splicing or 3′ end formation (Fig. 1E; Supplemental Fig. S1C). The rather low magnitude of these changes might seem surprising, when considering that nine sites for two families of relatively highly expressed miRNAs were disrupted. Nonetheless, the low magnitude of this effect was consistent with the ≤2-fold effects previously observed in reporter assays and in pancreatic islets from *Zeb1^200M^* mice (Gregory et al. 2008; Korpal et al. 2008; Park et al. 2008; Title et al. 2018; Title et al. 2021).

### Loss of miR-200 regulation of *Zeb1* causes impaired female fertility

*Zeb1^200H^* mice appeared normal and healthy and were interbred to generate *Zeb1^200M^*mice. The resulting litters had normal numbers of *Zeb1^200M^* mice at weaning (Table S1) and during the course of maintaining the colony, we noted no fertility defects in adult *Zeb1^200M^* males. However, the adult *Zeb1^200M^* female mice were subfertile. Here, we focused on this sex-specific fertility phenotype.

When paired with C57Bl/6J males for 12 weeks, 9 of 11 *Zeb1^200M^* females failed to produce any litters (Fig. 2A). The *Zeb1^200M^* females that did become pregnant had fewer litters than did wild-type or *Zeb1^200H^*females (Fig. 2A), and *Zeb1^200M^* litters tended to be smaller (Fig. 2B). *Zeb1^200M^*females had normal body weights (Supplemental Fig. S2A) and robust estrous cyclicity (Fig. 2C,D), indicating that the subfertility was likely not due to abnormal development of the hypothalamic–pituitary–ovarian axis.

**Figure 2.**
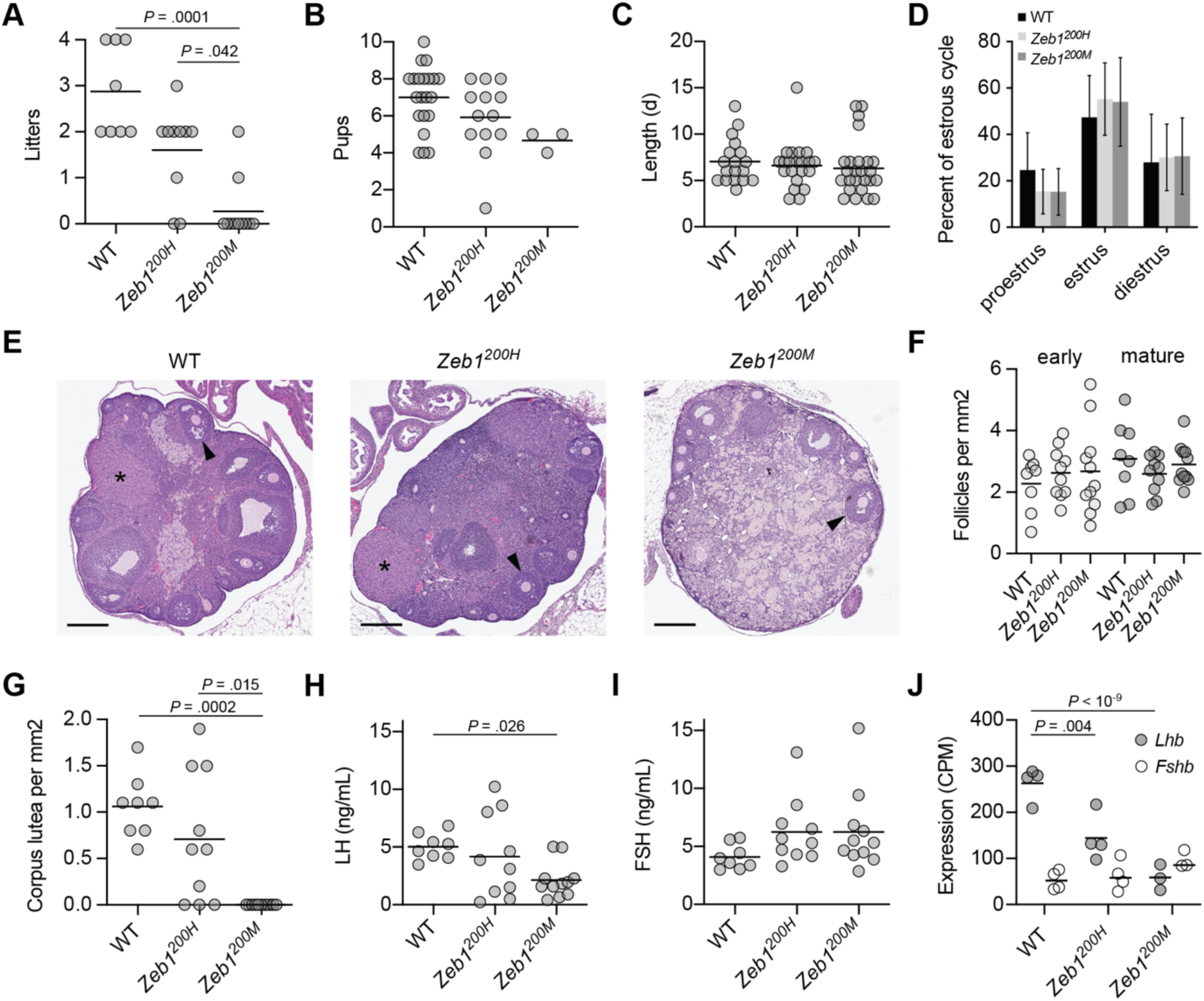
miR-200a/b-resistant *Zeb1* mice have decreased fertility. (A–B) Influence of *Zeb1^200^* alleles on number and size of litters. Plotted in (A) are the number of litters per female for WT, *Zeb1^200H^*, and *Zeb1^200M^* female mice mated with WT male mice for 12 weeks (black line, mean; n = 8–11 mice per genotype). Plotted in (B) are the number of pups per litter for the same cohort. Only statistically significant changes (adjusted *p* value <0.05) for all pairwise comparisons are indicated (ANOVA with Kruskal-Wallis test). Statistical testing for differences in litter size between *Zeb1^200M^*mice and the other two genotypes lacked sufficient power due to the small number of litters produced by *Zeb1^200M^* females. (C–D) Influence of *Zeb1^200^* alleles on estrus cycle length and stage. Plotted in (C) are the lengths in days for each estrus cycle, as determined by vaginal cytology, for WT, *Zeb1^200H^*, and *Zeb1^200M^*female mice (black line, mean; n = 18–23 estrus cycles from 8–11 mice per genotype). Plotted in (D) are the mean percentages of time (in days) spent in proestrus, estrus, and diestrus for the same cohort (error bars, s.d.). No statistically significant changes (adjusted *p* value <0.05) were detected (ANOVA with Kruskal-Wallis test). (E–G) Influence of *Zeb1^200^* alleles on ovarian folliculogenesis. Shown in (E) are hematoxylin and eosin stained sections of ovary from 27–40-week-old WT, *Zeb1^200H^*, and *Zeb1^200M^*female mice, post-breeding trial (arrowhead, secondary/mature follicle; asterisk, corpus luteum). The number of early/primary and mature/secondary follicles per mm2 ovary section and the number of corpus luteum per mm2 ovary section are plotted in (F) and (G), respectively, for the indicated genotypes (black line, mean; n = 8–11 mice per genotype). Only statistically significant changes (adjusted *p* value <0.05) for all pairwise comparisons are indicated (ANOVA with Kruskal-Wallis test). (H–I) Influence of *Zeb1^200^*alleles on circulating LH and FSH levels. Plotted are concentrations of LH (H) and FSH (I) in serum harvested from 27–40-week-old WT, *Zeb1^200H^*, and *Zeb1^200M^* female mice (black line, mean; n = 8– 11 mice per genotype) during pro-estrus, as determined by vaginal cytology. Only statistically significant changes (adjusted *p* value <0.05) for all pairwise comparisons are indicated (ANOVA with Kruskal-Wallis test). (J) Influence of *Zeb1^200^* alleles on *Lhb* and *Fshb* expression in the pituitary. Plotted are the counts per million mapped reads (CPM) for *Lhb* and *Fshb*, as determined by RNA sequencing, in WT, *Zeb1^200H^*, and *Zeb1^200M^*pituitary (black line, mean; n = 3–4 per genotype). Benjamini-Hochberg adjusted *p* values, as determined by a Wald test (DESeq2), are shown for *p* values less than 0.05.

To identify the basis of the impaired fertility in *Zeb1^200M^* females, we examined the ovaries of female mice in proestrus after the conclusion of the breeding trial. Grossly, ovaries were similar in size between different genotypes, and histology of *Zeb1^200M^*ovaries revealed follicles at all stages of development and similar numbers of follicles as wild-type and *Zeb1^200H^* ovaries (Fig. 2E,F). However, the number of corpus lutea was greatly diminished (Fig. 2G). Because corpus lutea formation depends on ovulation, the paucity of corpus lutea in *Zeb1^200M^* ovaries suggested ineffective ovulation.

The reduction in corpus lutea in *Zeb1^200M^* ovaries raised the possibility that LH synthesis and/or secretion might be dysregulated. We measured serum levels of LH and FSH during proestrus when LH levels typically surge. Indeed, circulating LH levels were decreased 2.4-fold in *Zeb1^200M^* females, whereas circulating FSH levels were not significantly changed (Fig. 2H,I). *Zeb1^200H^* females, which are fertile, had normal levels of both LH and FSH. To determine whether the decreased circulating LH might be caused by decreased LH production in the pituitary, we examined *Lhb* expression in our pituitary RNA-seq data. *Lhb* levels were reduced 1.8-fold in *Zeb1^200H^*pituitary and 4.5-fold in *Zeb1^200M^* pituitary (Fig. 2J), consistent with previous reports showing that the *Lhb* promoter, which contains at least four E-box motifs, is bound and repressed by ZEB1 (Hasuwa et al. 2013; Schultz et al. 2024).

In theory, the decreased expression of *Lhb* in *Zeb1* mutant pituitaries could be explained by either a reduction in the number of gonadotrope cells or a reduction in *Lhb* expression within the gonadotrope lineage. *Fshb*, which encodes the unique beta-subunit of follicle stimulating hormone, is also specifically expressed in gonadotropes. Unlike *Lhb*, *Fshb* levels were not affected in *Zeb1* mutants (Fig. 2J), indicating that the reduction in *Lhb* was due to reduced *Lhb* expression within the gonadotrope lineage and not a loss of gonadotropes.

### Massive gene expression changes accompany modest derepression of *Zeb1*

To gain a better understanding of the impact of elevated *Zeb1* on pituitary gene expression, we performed differential expression analysis using DESeq2 analysis (Love et al. 2014) of RNA-seq data from female mice that completed the pregnancy trial (Supplemental Table S2). In *Zeb1^200M^*pituitary, 2325 genes were differentially expressed (adjusted *p* value < 0.05), 43% of which were downregulated (Fig. 3A); in *Zeb1^200H^* pituitary, 1609 genes were differentially expressed, 43% of which were downregulated (Fig. 3B). In contrast, only 83 genes were differentially expressed when comparing *Zeb1^200H^*to *Zeb1^200M^* pituitary (Fig. 3C). The fold changes observed in *Zeb1^200H^* and *Zeb1^200M^* pituitary when compared to wild-type pituitary were highly correlated (Pearson *R*^2^ = 0.78) and linear regression of these changes yielded a slope of 1.08 (95% confidence interval 1.07–1.09), indicating a common dysregulated program of gene expression in the two genotypes with only slightly greater magnitude of change in the *Zeb1^200M^* pituitary (Fig. 3D). The similar changes observed for *Zeb1^200H^* and *Zeb1^200M^*pituitary indicated that these changes were not due to a difference in pregnancy history, as these two genotypes had divergent pregnancy outcomes. Consistent with these findings, principal component analysis (PCA) of the 2000 most variable genes also indicated that the gene expression profiles of *Zeb1^200H^*and *Zeb1^200M^* pituitaries were more similar to each other than to the profile of wild-type pituitary (Supplemental Fig. S3A). As described below, we attribute these striking similarities between the transcriptomes of the *Zeb1^200H^* and *Zeb1^200M^* pituitaries to the miR-200a/b–ZEB1/2 DNFL.

**Figure 3.**
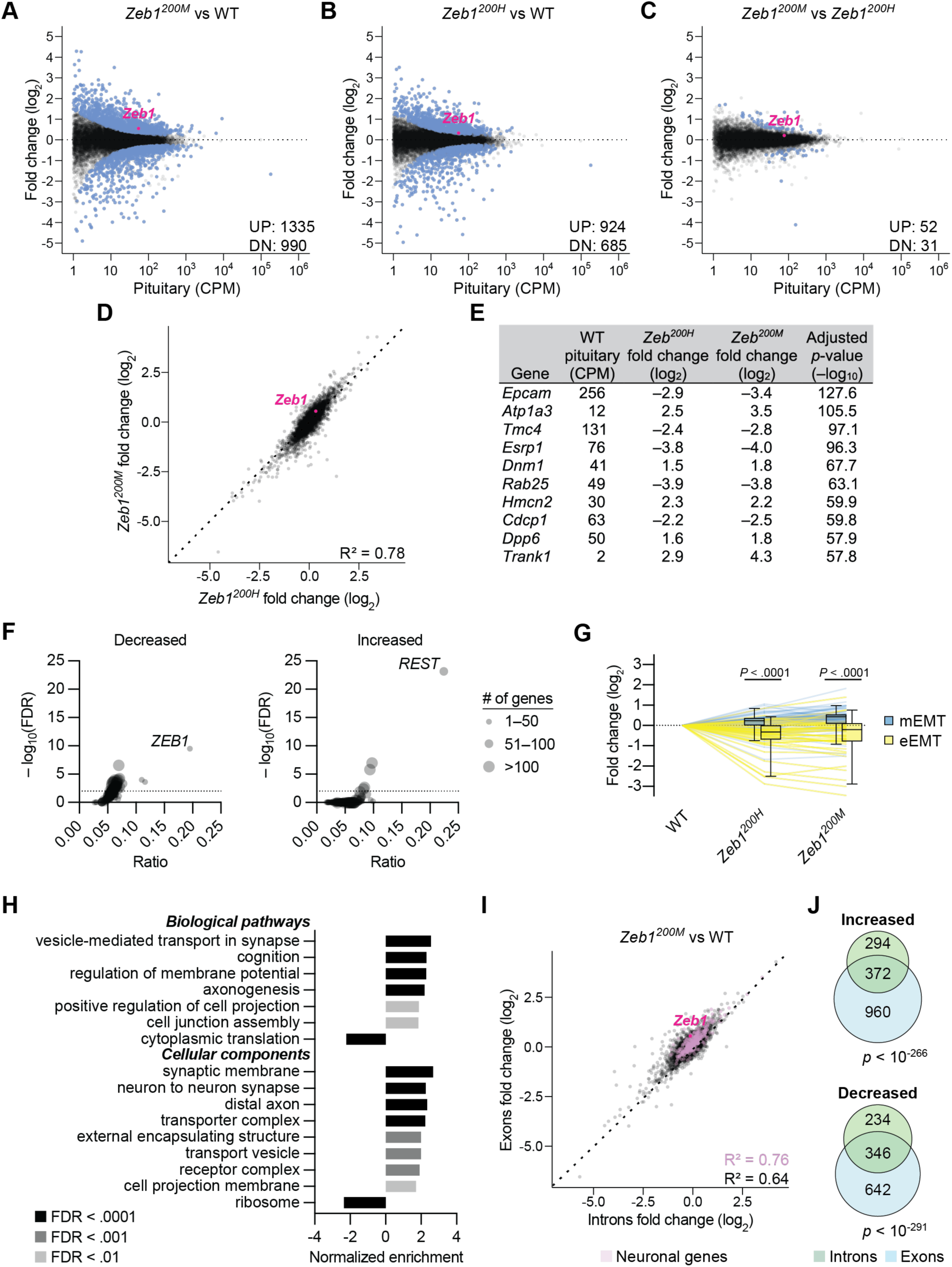
Widespread gene expression changes induced by modest derepression of *Zeb1*. (A–C) The influence of *Zeb1^200^*alleles on RNA levels in the pituitary. Shown are fold changes in mean RNA levels for *Zeb1^200M^* relative to WT (A), *Zeb1^200H^*relative to WT (B), and *Zeb1^200M^* relative to *Zeb1^200H^*(C), as determined by RNA sequencing and plotted as a function of expression in WT pituitary (n = 3–4 per genotype). Each circle represents a unique mRNA or noncoding RNA, showing results for all RNAs expressed above 1 count per million mapped reads (CPM). Blue circles indicate differentially expressed genes as determined by DESeq2 (adjusted *p* value < 0.05) and the number of significantly increased (UP) and decreased (DN) genes is indicated. (D) Correlation of RNA fold changes between *Zeb1^200^* alleles. Plotted are fold changes of *Zeb1^200M^* and *Zeb1^200H^*pituitary relative to WT pituitary. The correlation coefficient (Pearson *R*^2^) is indicated. (E) Top 10 differentially expressed genes, ranked by adjusted *p* value. For each gene, mean WT pituitary expression, and fold changes in *Zeb1^200H^* and *Zeb1^200M^* pituitary relative to WT pituitary are shown. (F) Overrepresentation of transcription factor binding sites in differentially expressed genes. Plotted are the ratio of genes with transcription factor binding sites that are also increased or decreased between *Zeb1^200M^* and WT pituitary (adjusted *p* value < 0.05) versus the false discovery rate, as determined by ChEA3. Each circle represents the high-confidence targets of an individual transcription factor assembled from ENCODE ChIP-Seq data. (G) The influence of *Zeb1^200^* alleles on genes associated with epithelial-mesenchymal transition. Plotted are the fold changes of 46 epithelial EMT (eEMT) genes and 62 mesenchymal EMT (mEMT) genes in *Zeb1^200M^*and *Zeb1^200H^* pituitary relative to WT pituitary. Statistical significance of the difference between eEMT genes and mEMT genes for each genotype are indicated (unpaired t-test). (H) Gene set enrichment analysis of *Zeb1^200M^* pituitary. Plotted are normalized enrichment scores for are all statistically significant (FDR < 0.01) biological pathway and cell component gene ontologies, as determined by WebGestalt (Elizarraras et al. 2024). (I) Correlation of RNA exon and intron fold changes in *Zeb1^200M^* pituitary. Plotted are exonic fold changes of *Zeb1^200M^* relative to WT pituitary versus intronic fold changes for the same comparison. The correlation coefficient (Pearson *R*^2^) is indicated, and neuronal genes are highlighted in lavender. (J) Overlap of differentially expressed genes (adjusted *p* value < 0.05) based on exonic read counts or intronic read counts. Shown are Venn diagrams for genes that are either increased or decreased in *Zeb1^200M^* relative to WT pituitary based on mapping to exons or introns. The significance of the overlap is indicated (Fisher’s exact test).

The modest but significant increase in *Zeb1* mRNA levels raised the question of whether ZEB1 targets might be affected. ZEB1 acts predominantly as a transcriptional repressor and some of its best-characterized targets are classic epithelial genes such as *Epcam* and *Cdh1*. Indeed, bona fide ZEB1 targets *Epcam*, *Esrp1*, and *Rab25* were among the most significantly differentially expressed genes, decreasing 7–15-fold in *Zeb1^200H^* pituitary and 11–16-fold in *Zeb1^200M^*pituitary (Fig. 3E).

To systematically characterize the contribution of ZEB1 and other transcription factors to the differential gene expression in *Zeb1^200M^*pituitary, we used ChEA3, a tool for analysis of enrichment of transcription-factor activity, which evaluates the overlap between user-provided gene lists and high-confidence targets for 118 transcription factors assembled from ENCODE ChIP-Seq data (Keenan et al. 2019). For each transcription factor, we plotted the gene ratio, which represents the fraction of genes in the ChIP-Seq set that were also differentially expressed in our RNA-seq data, versus the false discovery rate (FDR) (Fig. 3F). ZEB1 targets were the most overrepresented among the 990 genes that underwent decreased expression in the *Zeb1^200M^* pituitary (33 out of 169 targets, FDR < 10^−9^). ZEB1 targets were not overrepresented among the 1335 genes that underwent increased expression in *Zeb1^200M^*pituitary, indicating that in this tissue context ZEB1 is acting primarily as a transcriptional repressor rather than activator. Instead, we found that REST targets were the most overrepresented among the genes that underwent increased expression (89 out of 337 predicted targets, FDR < 10^−23^). REST is a transcriptional repressor that silences neuronal genes in non-neuronal cells (Jin et al. 2023). Unlike *Zeb1*, *Rest* mRNA was not differentially expressed in *Zeb1^200M^*pituitary, suggesting that its decreased activity was due to changes at the translational or post-translational level.

Given the central role of ZEB1 in EMT, we sought to determine whether EMT transcriptional changes were occurring in *Zeb1^200H^*and *Zeb1^200M^* pituitary. Using a core set of EMT genes derived from a meta-analysis of 18 independent studies (Gröger et al. 2012), we compared genes that are commonly downregulated during EMT (epithelial or eEMT genes) and genes that are commonly upregulated during EMT (mesenchymal or mEMT genes). In both *Zeb1^200H^* and *Zeb1^200M^* pituitary, eEMT genes were decreased, and mEMT genes were increased, with the magnitude of these effects somewhat greater in the *Zeb1^200M^* pituitary than in the *Zeb1^200H^* pituitary (Fig. 3G). Similar effects on eEMT and mEMT genes have been reported for *Zeb1^200M^* pancreatic beta-cells, indicating that these programs are largely independent of cell lineage (Title et al. 2018; Title et al. 2021).

To identify additional pathways affected by elevated pituitary *Zeb1*, we performed gene set enrichment (GSE) analysis. The most significantly depleted gene sets were related to ribosomal protein genes and translation, whereas the most significantly enriched gene sets were related to neuronal function and synaptic activity (Fig. 3H). Increased neuronal gene expression, combined with enrichment (i.e., derepression) of REST targets, suggested a model in which miR-200a/b-mediated repression of *Zeb1* enhances silencing of neuronal genes in ectoderm-derived anterior pituitary cells. Alternatively, increased neuronal gene expression could be secondary to changes in the posterior pituitary, which consists primarily of nerve endings from neurons that originate in the hypothalamus. In the former model, the gene-expression changes would affect both mature mRNAs and pre-mRNAs; in the latter model, the gene-expression changes would only affect mature mRNAs, as the pre-mRNAs are restricted to the hypothalamus. Therefore, we aligned our pituitary RNA sequencing data to intronic regions and performed differential expression analysis (Supplemental Table S3). A strong correlation between the fold changes of exonic and intronic regions was observed for all genes (Pearson *R*^2^ = 0.64) and for neuronal genes (Pearson *R*^2^ = 0.76) when comparing *Zeb1^200M^* to wild-type pituitary (Fig. 3I). We also compared the lists of significantly differentially expressed genes (adjusted *p* value < 0.05) from analysis of exonic and intronic regions and noted a statistically significant overlap for both upregulated and downregulated genes (Fig. 3J). As expected, ZEB1 targets featured prominently in the 346 genes with decreased expression of both exons and introns; of the 33 ZEB1 targets with decreased expression in *Zeb1^200M^* pituitary (Fig. 3F), 23 also had significantly decreased intronic expression. These findings support the hypothesis that the gene expression changes in the *Zeb1^200M^* pituitary are largely due to transcriptional changes in the anterior pituitary. Of note, *Zeb1* expression is increased in the *Zeb1* mutant pituitary when considering sequencing reads mapping to exons, but *Zeb1* expression is unchanged when considering intronic regions—the expected result, given that the mutant mouse model disrupts repression of *Zeb1* by the miR-200 families, which occurs post-transcriptionally.

### No evidence is found for *Zeb1* dysregulation in hypothalamus and ovary

As the miR-200a/b–ZEB1/2 axis has been implicated in hypothalamic control of sexual maturation and fertility (Messina et al. 2016), we also investigated gene expression in the hypothalamus of female *Zeb1^200H^* and *Zeb1^200M^* mice at 25–30 weeks of age (Supplemental Table S2). Unlike in the pituitary, *Zeb1* levels did not increase in the *Zeb1* mutant hypothalamus. Moreover, in contrast to the pituitary, a small number of genes (5 in *Zeb1^200H^*and 361 in *Zeb1^200M^*; adjusted *p* value < 0.05) were differentially expressed (Supplemental Fig. S3B–D), and PCA of the 2000 most variable genes poorly separated the individual genotypes (Fig. S3A). Together, these results indicated that miR-200a/b miRNAs regulate *Zeb1* in this tissue only very weakly, if at all, at least at this age.

As in the hypothalamus, *Zeb1* levels were unchanged in the *Zeb1^200H^* and *Zeb1^200M^* ovary compared to wild-type ovary. However, unlike in the hypothalamus, many more genes were differentially expressed in the ovary; 1319 genes were differentially expressed (adjusted *p* value < 0.05) in the *Zeb1^200M^*ovary, of which 46% were downregulated, whereas only 21 genes were differentially expressed in the *Zeb1^200H^* ovary (Supplemental Fig. S3E–G; Supplemental Table S2). PCA demonstrated that *Zeb1^200M^*ovaries were the most dissimilar, whereas *Zeb1^200H^* and wild-type samples failed to separate (Supplemental Fig. S3A). The striking difference in *Zeb1^200M^* ovary gene expression was presumably due to *Zeb1^200M^* females ovulating less frequently or not at all. Among the most downregulated genes were LH-responsive genes (e.g. *Runx2*, *Sfrp4*), steroidogenic enzyme genes involved in placental progesterone synthesis (e.g. *Star*, *Cyp11a1*), and/or corpus luteum marker genes (Supplemental Fig. S3H). These findings are consistent with the ovarian dysfunction observed in *Zeb1^200M^* mice arising because of pituitary LH insufficiency.

### Increased *Zeb1* levels inhibit miR-200a/b expression

Given the well-known reciprocal repression that occurs between ZEB1 and miR-200a/b, we next sought to determine whether miR-200a/b levels were altered in *Zeb1^200H^* and *Zeb1^200M^* pituitary. First, we assessed expression of the primary transcripts that host the miR-200a/b miRNAs using our pituitary RNA-seq data. In wild-type pituitary, pri-miR-200b∼a∼429 levels were ∼3-fold higher than pri-miR-200c∼141 levels (Fig. 4A). In *Zeb1^200H^* and *Zeb1^200M^* pituitaries, pri-miR-200b∼a∼429 levels were decreased 5.7-fold and 7.7-fold, respectively, and pri-miR-200c∼141 levels were decreased 2.9-fold and 3.2-fold, respectively, consistent with previous reports demonstrating that miR-200 loci are direct targets of ZEB1 (Bracken et al. 2008; Burk et al. 2008; Balestrieri et al. 2018) (Fig. 4A).

**Figure 4.**
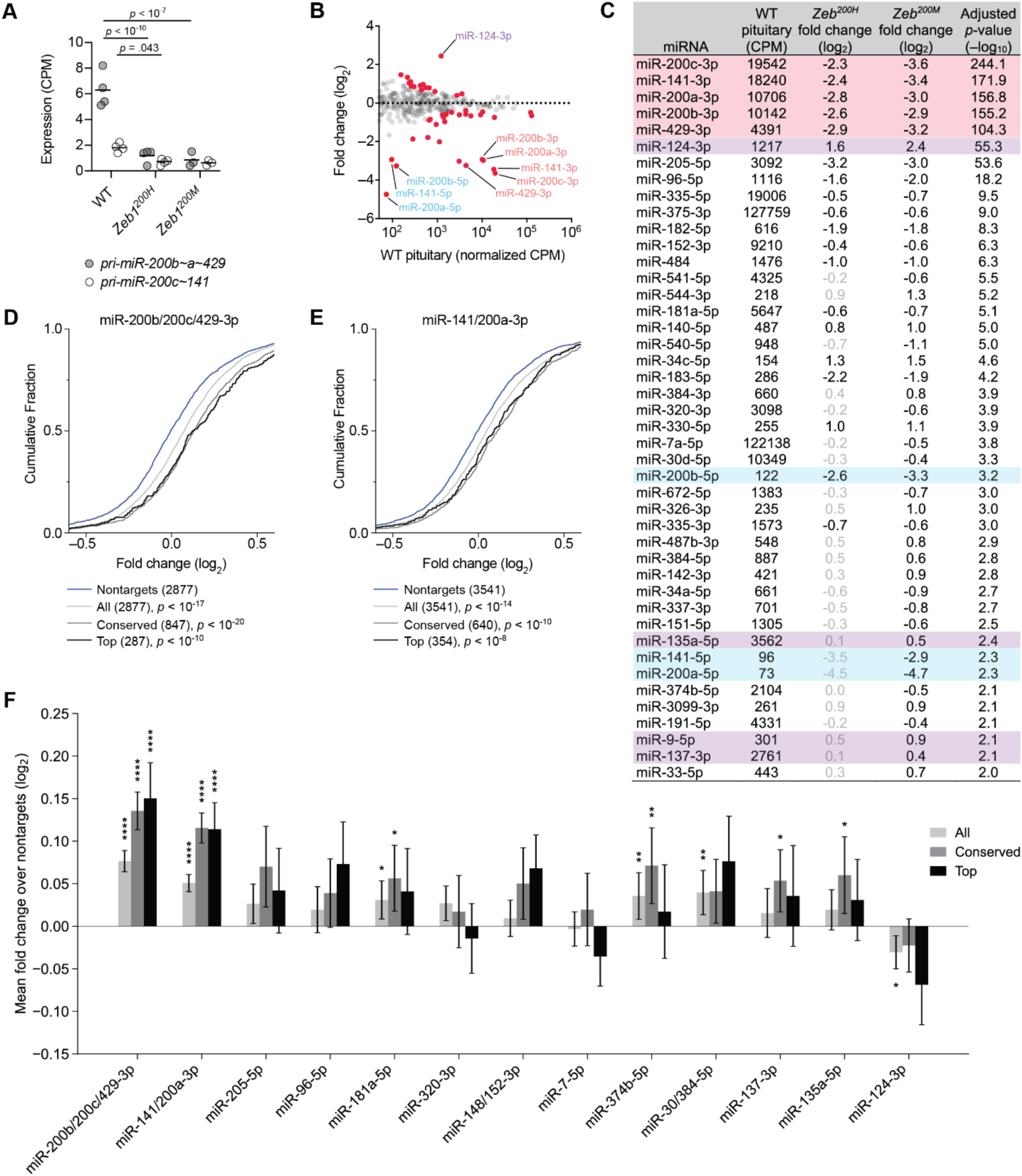
miR-200a/b-resistant *Zeb1* mice have decreased expression of miR-200a/b and increased expression of miR-200a/b targets in the pituitary. (A) Influence of *Zeb1^200^* alleles on miR-200a/b family primary transcripts in the pituitary. Plotted are the counts per million mapped reads (CPM) for *pri-miR-200b∼a∼429* and *pri-miR-200c∼141*, as determined by RNA sequencing, in wild-type, *Zeb1^200H^*, and *Zeb1^200M^*pituitary (black line, mean; n = 3–4 per genotype). Benjamini-Hochberg adjusted *p* values, as determined by a Wald test (DESeq2), are shown for *p* values less than 0.05. (B–C) The influence of *Zeb1^200M^* on mature miRNA levels in the pituitary. Plotted in (B) are fold changes in mean miRNA levels for *Zeb1^200M^* pituitary relative to wild-type pituitary (n = 3–4 per genotype), as determined by small-RNA sequencing and plotted as a function of expression in wild-type pituitary. Each circle represents one miRNA, showing results for all miRNAs expressed above 1 count per million mapped miRNA reads (CPM). Red circles indicate differentially expressed miRNAs (adjusted *p* value < 0.01). For each miRNA indicated by a red circle in (B), mean wild-type pituitary expression, fold changes in *Zeb1^200H^*and *Zeb1^200M^* pituitary relative to wild-type pituitary, and adjusted *p* value for the *Zeb1^200M^* comparison are shown in (C). Pink denotes miRNAs of the miR-200a/b families, blue denotes passenger strands of the miR-200a/b families, and lavender denotes brain-specific miRNAs. Fold changes indicated in light gray are not statistically significant (adjusted *p* value > 0.01). (D–E) The influence of *Zeb1^200M^*on expression of miR-200b/200c/429-3p and miR-141/200a-3p predicted targets. Plotted are the cumulative distribution functions of mRNA fold changes in the *Zeb1^200M^* pituitary relative to wild-type pituitary for all predicted targets (All; light grey), conserved predicted targets (Conserved; grey), top 10% predicted targets (Top; black). The number of genes in each target set is indicated. *P* values were calculated based on a Mann-Whitney test between each target set and a set of 3′ UTR length-matched transcripts with no predicted site for each miRNA family randomly sampled at a 1:1 ratio. For simplicity, only the nontargets matched to the all targets set are plotted (blue). Selection of nontarget cohorts for each target set was repeated 21 times, and the iteration generating the median *p* value is displayed. (F) The influence of *Zeb1^200M^* on expression of predicted targets of differentially expressed miRNAs. Plotted are the differences between the median fold change of each set of predicted targets (all, conserved, top) and the median fold change of its matched set of nontargets. This metric of repression was calculated for 21 sampled sets of nontargets, and the mean is displayed (error bars, standard deviation). For each sampling, a *p* value was calculated as described in (D–E) and the median *p* value is displayed (*, *p* < 0.05; **, *p* < 0.01, ***, *p* < 0.001, ****, *p* < 0.0001). miR-200 family targets were excluded when calculating target repression for other miRNA families. Results for differentially expressed miRNAs with at least 1000 CPM in wild-type and at least 100 conserved targets (after removing miR-200 family targets) are plotted.

Small-RNA sequencing of pituitaries identified 44 and 20 miRNAs that were differentially expressed in *Zeb1^200M^*and *Zeb1^200H^* pituitary, respectively (Fig. 4B,C; Supplemental Fig. S4A; Supplemental Table S4). The most significant of these were miR-200a/b family miRNAs, which were decreased 5.0–7.4-fold in *Zeb1^200H^*and 7.7–12.5-fold in *Zeb1^200M^* pituitary (Fig. 4B,C, pink). Consistent with the decreased levels of primary transcript, miR-200a/b passenger strands meeting the expression cut-off in wild-type pituitary were also substantially reduced (Fig. 4B,C, blue). In addition, several miRNAs had increased expression in *Zeb1* mutant pituitaries, including the brain-specific miRNAs miR-124-3p, miR-135a-5p, miR-9-5p, and miR-137-3p (Fig. 4B,C, lavender). Similar to the mRNA fold changes, the miRNA fold changes observed in *Zeb1^200H^* and *Zeb1^200M^* pituitary were highly correlated (Pearson *R*^2^ = 0.82), and linear regression of these changes yielded a slope of 1.043 (95% confidence interval 0.986–1.101), indicating a common dysregulated program of gene expression in the two genotypes (Supplemental Fig. 4B).

The striking reduction in miR-200a/b levels in *Zeb1* mutant pituitaries raised the question of whether miR-200a/b targets might be affected. To answer this question, we compared fold changes of predicted miR-200a or miR-200b targets to fold changes of mRNAs lacking a miR-200a/b site. The effects on all predicted targets, conserved predicted targets, and the top 10% of predicted targets (Agarwal et al. 2015) were compared to UTR length-matched control sets with the expectation that higher confidence predictions should correlate with larger effects. Indeed, we observed increased levels of both miR-200a and miR-200b targets in *Zeb1^200M^* pituitary (Fig. 4D– F) and *Zeb1^200H^* pituitary (Supplemental Fig. S4C–E).

One of the top predicted miR-200a/b targets is *Zeb2* (Fig. 1B). Indeed, *Zeb2* mRNA levels increased 1.5-fold in *Zeb1^200H^*and 1.4-fold in *Zeb1^200M^* pituitary. Although these increases were not considered significant after adjustment for multiple hypotheses (Fig. 1E, *p* = 0.08 for *Zeb1^200H^* and *p* = 0.26 for *Zeb1^200M^*), they were significant after aggregating the data and comparing WT with the combined mutant data (*p* = 0.012 when pooling data for *Zeb1^200H^* and *Zeb1^200M^*). Because *Zeb2* is a close paralogue of *Zeb1*, these increases are expected to reinforce the effects of disrupting *Zeb1* repression. A similar reinforcement would be expected for the wild-type allele of *Zeb1^200H^*animals, as derepression of the mutant allele causes increased ZEB1 activity, which reduces miR-200a/b expression and miR-200a/b-mediated repression of wild-type *Zeb1* transcripts. Indeed, we found similar numbers of RNA-seq reads overlapping wild-type and mutant miR-200a/b sites in in *Zeb1^200H^* pituitary (Supplemental Fig. S4F), indicating that the DNFL nearly fully abrogates miR-200a/b repression of wild-type Zeb1 allele. In fact, the substantially higher expression of *Zeb1* than *Zeb2* observed in pituitary of wild-type animals (Fig. 1E) suggests that in *Zeb1^200H^* animals, derepression of the *Zeb1* wild-type allele has a greater effect than derepression of both *Zeb2* alleles.

Evidence of altered miRNA-mediated repression was not limited to miR-200a/b targets (Fig. 4F). Predicted targets of several other downregulated miRNAs also increased, although the highest-confidence predictions did not always display the greatest magnitude of change. Predicted targets of upregulated miRNAs did not decrease significantly, perhaps due to the generally lower abundance of these miRNAs. When considering the dozens of differentially expressed miRNAs in the *Zeb1^200H^* and *Zeb1^200M^* pituitary, miR-200a and miR-200b family members exerted the greatest effect on target repression, consistent with these miRNAs being both highly expressed and most significantly depleted in mutant pituitary.

These results, in concert with analysis of ZEB1 transcriptional targets, indicated that disrupting the miR-200a/b–ZEB1/2 DNFL directly influences expression of dozens if not hundreds of genes. These primary effects occurred through enhanced transcriptional repression by ZEB1 and reduced post-transcriptional repression by miR-200a/b. When adding on downstream, secondary effects, expression of more than 2000 genes was affected.

## Discussion

In this study, we characterize the physiological consequences of derepressing a single miRNA target in mice. Loss of miR-200a/b regulation of *Zeb1* caused anovulatory infertility and reduced LH expression in female mice. Some previous studies have also mutated miRNA complementary sites to investigate the phenotypic consequences of disrupting miRNA regulation of a single target (Lai and Posakony 1997; Dorsett et al. 2008; Teng et al. 2008; Cassidy et al. 2013; Ecsedi et al. 2015; Lu et al. 2015; Drexel et al. 2016; McJunkin and Ambros 2017; Mildner et al. 2017; Aeschimann et al. 2019; Garaulet et al. 2020; Yang and McJunkin 2020; Yang et al. 2020; Kuzniewska, 2022 #552; Young et al. 2022; Hurtado et al. 2024; Janati-Idrissi et al. 2024). In mice, this approach has revealed that miRNA regulation of a single gene can improve synaptic plasticity and memory (Kuzniewska et al. 2022), alter bone development (Young et al. 2022), and affect various phenomenon in immune cells, including response to infection (Lu et al. 2015), proliferation of thymocytes (Mildner et al. 2017), and derepression of cytidine deaminase activity (Dorsett et al. 2008; Teng et al. 2008). With respect to fitness of the organism, the fertility phenotypes we observed upon disruption of miR-200a/b repression of murine *Zeb1* are the most severe phenotypes observed for reduced miRNA regulation of a single mammalian mRNA and are among the most severe observed for mutation of a single mammalian 3′ UTR.

The fertility defects of mice with this miR-200a/b-resistant *Zeb1* 3′ UTR mirrored the fertility defects previously reported for miR-200b/429 double-knockout mice (Hasuwa et al. 2013) and for mice with gonadotrope-specific overexpression of ZEB1 (Hasuwa et al. 2013). Our results, combined with the results of these previous studies, converge on a model in which derepression of *Zeb1* is the driving cause of infertility associated with loss of miR-200b/429, and more generally, that miR-200a/b limits *Zeb1* expression in pituitary gonadotrope cells to promote *Lhb* expression and ovulation (Fig. 5).

**Figure 5.**
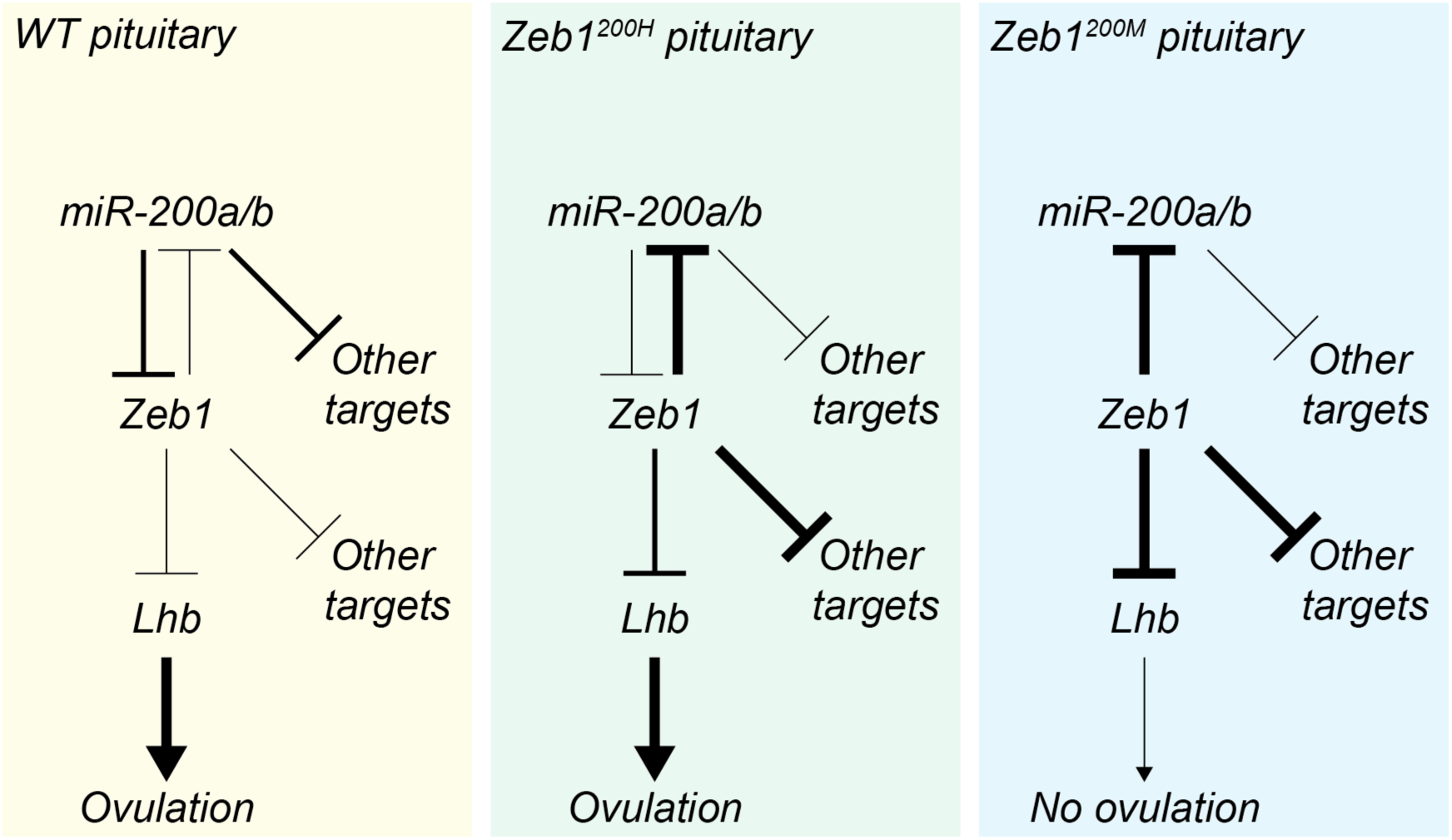
Increased ZEB1 activity in *Zeb1^200H^* and *Zeb1^200M^*pituitary represses miR-200a/b miRNAs, *Lhb*, and other ZEB1 targets. Line thicknesses indicate relative strengths of repression. Only *Zeb1^200M^* mice have enough ZEB1 activity to impair ovulation.

### Primary defect occurs in the pituitary

Although *Zeb1^200M^* mice are resistant to miR-200a/b in all cells and tissues, derepression of *Zeb1* within the hypothalamic-pituitary-ovarian axis appears to be restricted to the pituitary. Increased *Zeb1* expression was detected in *Zeb1^200M^* pituitary but not in *Zeb1^200M^*hypothalamus or ovary (Fig. 1E; Supplemental Fig. S3B,E). Likewise, increased repression of well-established ZEB1 targets, such as *Epcam*, *Esrp1*, and the miR-200a/b family primary transcripts, was observed exclusively in pituitary (Fig. 3E; Fig. 4A). These results are consistent with previous work demonstrating that ZEB1 protein is increased in the pituitary but not the hypothalamus of miR-200b/429 double-knockout mice and that forced expression of ZEB1 specifically in gonadotrope cells recapitulates the anovulation observed in miR-200b/429 double knockout mice (Hasuwa et al. 2013).

On the other hand, we found no evidence for a role of the miR-200a/b–Zeb1 DNFL in hypothalamus. Hypothalamic neurons produce gonadotropin-releasing hormone (GnRH), which stimulates expression and secretion of LH and FSH in the anterior pituitary. Complete loss of miRNAs in hypothalamic neurons blunts GnRH expression and sexual maturation, which ultimately results in both male and female sterility (Messina et al. 2016). Although miR-200a/b regulation of *Zeb1* is reported to mediate hypothalamic control of GnRH expression (Messina et al. 2016), our miR-200a/b-resistant *Zeb1* mice had normal sexual maturation; male *Zeb1^200M^* mice were fertile, and female *Zeb1^200M^* mice had normal estrous cycles (Fig. 2D; Supplemental Fig. S2B) and normal-sized ovaries. Thus, miR-200a/b regulation of *Zeb1* is not required for proper hypothalamic control of GnRH expression.

### *Zeb1* levels are tightly regulated

The 1.5-fold increase in *Zeb1* transcripts in *Zeb1^200M^* pituitary was modest but in line with previous observations. Similar *Zeb1* fold-changes are reported in cancer cell lines with reduced levels of miR-200a/b and in *Zeb1^200M^* pancreatic islets (Gregory et al. 2008; Korpal et al. 2008; Park et al. 2008; Title et al. 2018; Title et al. 2021). As in the pituitary, these fold-changes have functional consequences. For example, the ∼1.5-fold increase in *Zeb1* mRNA in pancreatic islets, accompanied by a ∼1.3-fold increase in ZEB1 protein, is sufficient to promote EMT, tumor invasion, and increased metastasis in two different autochthonous cancer models (Title et al. 2018).

How does the magnitude of *Zeb1* derepression compare to that of other miRNA targets? In general, median fold-changes of derepression in miRNA knockout and knockdown experiments range from ∼1.1–1.3-fold for sets of top predicted targets, with some of the variation in observed fold-changes presumably due to differences in miRNA abundance and/or incomplete depletion of the miRNA (Agarwal et al. 2015; Ahmed et al. 2017). Of course, most mRNAs that are targeted by one miRNA are also targeted by other miRNAs, and thus the overall effect of miRNAs on protein output can be much higher than 1.3-fold (Friedman et al. 2009). In *Zeb1^200M^* pituitary, which has ∼10-fold lower expression of miR-200a and miR-200b families than WT pituitary (Fig. 4B), median fold changes of derepression were ∼1.1-fold for top predicted miR-200a/b targets (Fig. 4F). In that regard, the 1.5-fold increase in *Zeb1* was well above the median. Indeed, both *Zeb1* and *Zeb2* ranked among the 20% most derepressed targets within our set of predicted top targets. We also cannot exclude the possibility that ZEB1 protein levels were further increased due to derepression of translation in miR-200a/b-resistant *Zeb1* pituitary.

Another way we might have underestimated the magnitude of derepression was if miR-200a/b regulation of *Zeb1* occurs in only some of the ∼12 distinct pituitary cell types. However, our data suggest that increased ZEB1 activity was likely occurring throughout the pituitary. In single-nuclei RNA-seq from WT female mouse pituitary, *Zeb1* is expressed at similar levels across the six main hormone-producing cell types, which account for more than 80% of all cells in the pituitary (Ruf-Zamojski et al. 2021). Likewise, ZEB1 targets *Epcam*, *Esrp1*, and *Rab25,* which decreased 11–16-fold in *Zeb1^200M^* pituitary, are each expressed at similar levels across the hormone-producing cell types (Ruf-Zamojski et al. 2021). Because these ZEB1 targets are broadly expressed at similar levels, the striking fold changes we observed in *Zeb1^200M^* pituitary were only possible if ZEB1 activity increased in most pituitary cells.

### *Zeb1* is dosage-sensitive

To our surprise, small changes in *Zeb1* expression were associated with a broad remodeling of the pituitary transcriptome (Fig. 3A). Some of the best-characterized ZEB1 targets, including *Epcam*, *Esrp1*, and the two miR-200a/b primary transcripts, were reduced 3–16-fold in *Zeb1^200M^* pituitary, suggesting very high sensitivity to *Zeb1* dosage (Fig. 3E; Fig. 4A). The mechanistic basis for this sensitivity to ZEB1 dosage is not known, however, we speculate that *cis*-acting features associated with ZEB1 binding sites, such as the number of CAGGTR motifs, proximity of binding sites for other transcription factors, and chromatin accessibility, might determine whether a ZEB1 target is sensitive to or buffered against modest changes in *Zeb1* expression, as has been reported for other transcription factors (Hannon et al. 2017; Naqvi et al. 2023).

The gene expression profiles of *Zeb1^200H^*and *Zeb1^200M^* pituitary were highly correlated, with only slightly larger-magnitude fold changes in *Zeb1^200M^* pituitary when each was compared to WT pituitary (Fig. 3D). For instance, in *Zeb1^200H^* pituitary, *Zeb1* expression increased to ∼86% of the level observed in *Zeb1^200M^* pituitary (Fig. 1E; Fig. 3A,B) and ZEB1 targets *Epcam*, *Esrp1*, *Rab25,* and the two miR-200a/b primary transcripts decreased in *Zeb1^200H^* pituitary by 67–105% of the amount observed in *Zeb1^200M^* pituitary (Fig. 3E; Fig. 4A). We attribute these transcriptomic similarities between the *Zeb1^200H^* and the *Zeb1^200M^* pituitary to the DNFL, in which derepression of the mutant *Zeb1^200H^* allele causes increased ZEB1 activity, which reduces *Mir200* transcription and miR-200a/b-mediated repression of mRNA from the wild-type *Zeb1^200H^* allele, such that the combined level of both *Zeb1* alleles of the *Zeb1^200H^* pituitary resembles that of the *Zeb1^200M^*pituitary.

Despite the high degree of overlap between *Zeb1^200H^*and *Zeb1^200M^* pituitary at the molecular level, only *Zeb1^200M^* mice had ovulation defects and impaired fertility (Fig. 2A,G). Two lines of evidence indicate that this phenotypic difference was due to differences in LH expression observed between *Zeb1^200H^*and *Zeb1^200M^* mice. First, administration of hormones used for superovulation restores ovulation in miR-200b/429 double knockout mice (Hasuwa et al. 2013). Second, *Lhb* transcripts, *Lhb* intron-mapping reads, and circulating LH protein are each decreased about 50% less in *Zeb1^200H^* mice than in *Zeb1^200M^* mice (Fig. 2H,J; Supplemental Table S3). This proposed mechanism implies that *Lhb* transcription is exquisitely sensitive to the small increase in *Zeb1* levels observed between *Zeb1^200H^* and *Zeb1^200M^* pituitary. The average difference in *Zeb1* levels, which we measured to be 15%, was so small that it did not achieve statistical significance after adjusting for multiple hypothesis testing in the RNA-seq analysis comparing *Zeb1^200H^*and *Zeb1^200M^* pituitary. Added translational derepression might accentuate this 15% difference, however, the effects are likely modest given the overall similarity in gene expression profiles between *Zeb1^200H^*and *Zeb1^200M^* pituitary (Fig. 3D). Thus, we infer that some small and likely difficult-to-quantify increase in ZEB1 between *Zeb1^200H^*and *Zeb1^200M^* pituitary must somehow explain the sizable *Lhb* decrease and fertility phenotype. One possibility is that transcription of *Lhb* is ultra-sensitive to the level of ZEB1, perhaps through highly cooperative ZEB1 binding to multiple sites in the *Lhb* promoter (Hasuwa et al. 2013; Schultz et al. 2024), such that a small increase in ZEB1 when comparing *Zeb1^200M^*to *Zeb1^200H^* pituitary is sufficient to reduce *Lhb* transcription more than 2-fold. Alternatively, the genes that undergo differential expression in *Zeb1^200M^* pituitary compared to *Zeb1^200H^* pituitary might include multiple regulators that cooperatively converge with ZEB1 to reduce *Lhb* transcription.

### Importance of the miR-200a/b–ZEB1/2 DNFL

Our study of miR-200a/b-resistant *Zeb1* mice, together with the previously reported findings of miR-200b/429 double-knockout mice (Hasuwa et al. 2013), demonstrates that miR-200a/b-mediated repression of *Zeb1* increases pituitary *Lhb* expression and circulating LH protein to physiological levels that support ovulation and female fertility (Fig. 5). This phenotype observed in miR-200a/b-resistant *Zeb1* mice differs from the phenotype observed with complete loss of *Lhb* expression, which causes hypogonadism and infertility in both male and female mice (Ma et al. 2004), indicating that the small amount of circulating LH detected in female miR-200a/b-resistant *Zeb1* mice is sufficient for gonadogenesis. Alternatively, because our measurements of *Lhb* transcripts and circulating LH protein were all taken from adult female mice during proestrus, when *Lhb* transcription and LH secretion are maximal, the molecular changes we observed might be limited to adult mice or to one or more stages of the estrus cycle. Age- and stage-resolved measurements of *Zeb1*, miR-200a/b, and *Lhb* in WT and *Zeb1^200M^*pituitary gonadotropes will be necessary to determine whether the miR-200a/b–ZEB1/2 DNFL acts continuously to suppress ZEB1 activity and promote *Lhb* transcription or serves a more dynamic purpose, for example, as an ultrasensitive, inducible switch for *Lhb* expression.

Like LH secretion, *Lhb* transcription is induced during proestrus by changes in the frequency of hypothalamic gonadotropin-releasing hormone (GnRH) pulses (Dalkin et al. 1989; Haisenleder et al. 1991; Kaiser et al. 1997; Thompson and Kaiser 2014). This GnRH signal is transduced by EGR1, SF-1, and PITX1, transcriptional activators that bind directly to the *Lhb* promoter (Halvorson et al. 1996; Halvorson et al. 1998; Tremblay and Drouin 1999; Wolfe and Call 1999). Less is known about the repressive factors that might restrict the duration and amplitude of GnRH-induced *Lhb* transcription. Our study and prior findings in mice and pituitary cell lines indicate that ZEB1 may play such a role: ZEB1 binds the *Lhb* promoter (Hasuwa et al. 2013; Schultz et al. 2024), and increased expression of ZEB1 blocks GnRH- or EGR1-stimulated *Lhb* expression in the murine LbetaT2 gonadotrope cell line (Schultz et al. 2024) and potently suppresses *Lhb* in vivo (Fig. 2J) (Hasuwa et al. 2013), whereas transient knockdown of *Zeb1* in LbetaT2 cells increases *Lhb* levels (Hasuwa et al. 2013).

Although increased ZEB1 activity in pituitary gonadotropes impairs female fertility, decreased ZEB1 activity appears to be better tolerated. Female mice with gonadotrope-specific deletion of *Zeb1* are fertile and have normal levels of pituitary *Lhb* and circulating LH, at least during diestrus (Schultz et al. 2024). Further investigation is needed to determine the impact of gonadotrope-specific ZEB1 deficiency on *Lhb* expression during other stages of the estrus cycle and after treatment with a GnRH agonist. Because constitutive overexpression of *Lhb* in gonadotropes causes anovulatory infertility and ovarian cysts and tumors (Risma et al. 1995), we speculate that mammals might have redundant mechanisms to limit *Lhb* expression and secretion.

Is ZEB1 regulation of Lhb conserved? To date, the effects of physiological levels of ZEB1 on *Lhb* expression has been reported only in mice and murine cell lines. When examining the effects of ZEB1 ectopic overexpression, a partial fragment of the human *LHB* promoter is detectably repressed, although the magnitude of repression is less than that observed with an orthologous fragment of the mouse *Lhb* promoter (Schultz et al. 2024). This differential sensitivity is partly attributed to two E-box motifs that are present in the mouse *Lhb* promoter but not in the human *LHB* promoter. The distal motif is also found in ungulates (pigs, cows) and elephants, and the proximal motif is found in most placental mammals surveyed, except for humans and chimpanzees (Supplemental Fig. S5A). For both mouse and human promoter fragments, some of the ZEB1 repression appears to be independent of canonical E-box motifs, suggesting either non-canonical binding or indirect effects of ZEB1. Also worth noting that the promoter fragments used in this study end shortly after the TSS and thus do not include an additional E-box motif downstream of the TSS (+18 bp), which is the most deeply conserved E-box motif within 500 bp of the TSS in either direction (Supplemental Fig. S5A). The importance of this motif in ZEB1-mediated repression of *Lhb* awaits further investigation.

Although *Lhb* is restricted to placental mammals, orthologues of both miR-200 and ZEB1/2 have existed for more than 600 million years, i.e., since the last common ancestor of Bilateria. Even so, the miR-200–ZEB DNFL appears to be a more recent innovation. Multiz alignments of the *Zeb1* and *Zeb2* 3′ UTRs indicate that miR-200a/b repression of *Zeb1* and *Zeb2* likely originated with vertebrates (Fig. 1C, Supplemental Fig. S1C). Multiz alignments of the *Mir200b∼a∼429* and *Mir200c∼141* promoters suggest that ZEB1/2 repression of *Mir200b∼a∼429* also likely originated with vertebrates, whereas ZEB1/2 repression of the mammalian-specific *Mir200c∼141* cluster likely originated with placental mammals (Supplemental Fig. S5C). While most functional studies of the miR-200a/b–ZEB1/2 have focused on mice and human cell lines, one group reported that overexpression or morpholino-based knockdown of *zeb1a/b* in zebrafish causes a ∼2-fold decrease or increase in miR-200a/b miRNAs, respectively (Vannier et al. 2013). Nevertheless, miR-200a/b knockdown did not recapitulate the developmental cell adhesion defects observed with overexpression of *zeb1a/b*, indicating that the miR-200a/b–ZEB1/2 DNFL is likely not required for the developmental function ascribed to *zeb1a/b*. Understanding how the miR-200a/b– ZEB1/2 DNFL is used in zebrafish and other animals may reveal new and common principles about this ancient gene regulatory circuit.

## MATERIALS AND METHODS

### Mouse husbandry

Mice were group-housed in a 12-hr light/dark cycle (light between 07:00 and 19:00) in a temperature-controlled room (21.1 ± 1.1°C) at the Whitehead Institute for Biomedical Research with free access to water and food and maintained according to protocols approved by the Massachusetts Institute of Technology Committee on Animal Care. Euthanasia was performed by CO_2_ inhalation.

### Generation of *Zeb1^200^* and *Zeb2^200^* mice

The previously reported *Zeb1^200H^* knock-in mice (Title et al. 2018) were backcrossed to C57BL/6J mice for at least 15 generations prior to initiating survival and fertility studies. The survival of heterozygous (*Zeb1^200H^*) and homozygous (*Zeb1^200M^*) knock-in mice was confirmed in mouse lines derived from the two independent ES cell clones (6B6 and 3C5) (Table S1). The *Zeb1^200H^* knock-in mice derived from clone 6B6 have been deposited at JAX with the MGI ID Zeb1<TM1DPBL>.

Heterozygous *Zeb2^200H^* knock-in mice were generated as follows. To generate the targeting construct, a DNA fragment containing the mouse *Zeb2* locus was subcloned from the bacterial artificial chromosome RP23-454M20. The targeting arms spanned last exon of *Zeb2*, which includes the entire 3′ UTR (Supplemental Fig. S1C). All eleven miR-200a/b sites were mutated by site-directed mutagenesis. A LoxP-flanked puromycin-resistance gene was used for positive selection and diphtheria toxin gene (DTA) was used for negative selection. The targeting construct was electroporated into v6.5 mouse embryonic stem cells (genotype 129SvJae x C57Bl/6; male) and 236 colonies were screened by PCR and Southern blot (Supplemental Table S5), yielding 21 positive clones. Two clones with the first nine miR-200a/b sites mutated (1D2 and 2A4) and one clone with first ten sites mutated (3E3) were injected into C57BL/6J blastocysts and transferred into pseudo-pregnant females. Chimeras were bred with C57BL/6J mice to generate heterozygous progeny and germline transmission of the mutant allele was verified by PCR. The puromycin resistance cassette was excised by intercrossing with transgenic CMV-cre mice. Heterozygous knock-in mice were backcrossed to C57BL/6J mice for at least 10 generations. The *Zeb2^200H^*mice derived from clone 3E3 (first 10 sites mutated) and clone 1D2 (first 9 sites mutated) have been deposited at JAX with the MGI IDs Zeb2<TM1DPBL> and Zeb2<TM2DPBL>, respectively.

### Genotyping

Genomic DNA was extracted from mouse earsnips using the HotSHOT method (Truett et al. 2000). PCR was performed using a two-step protocol (98°C for 10s, 72°C for 10s) with 1 μL of genomic DNA, Phusion Flash High-Fidelity PCR Master Mix (Thermo Scientific), and 10 pmoles of primers (Supplemental Table S5) and then analyzed by agarose gel electrophoresis.

### Experimental design and breeding trial

A cohort of WT (n=9), heterozygous *Zeb1^200H^* (n=10), and homozygous *Zeb1^200M^* (n=11) female mice were generated using both clonal lines (17 animals from 3C5, 11 animals from 6B6/3C5 intercross, and 2 animals from 6B6) and singly housed after weaning and genotyping. Starting at 8–11 weeks of age, mice were subjected to 3 weeks of estrous cycle staging by vaginal cytology. Of the 30 mice in our cohort, 29 completed at least one estrus cycle and were included in the subsequent breeding trial. One WT mouse never cycled and was excluded from further studies. Breeding was initiated by adding a WT C57Bl/6J male mouse to each cage. During the 12-week breeding trial, animals were assessed daily for visible pregnancies and new litters. For half of the mice in the trial, including all *Zeb1^200M^* females, breeding was extended an additional 5 weeks although no additional litters were produced by *Zeb1^200M^* females. Only the results from the first 12 weeks of breeding are plotted in Figure 2A. At the conclusion of the trial, male mice were removed from cages and, after 20 or more days, females were again staged for estrous cycle. At the first instance of proestrus, female mice (27–40 weeks of age) were euthanized by CO2 inhalation, blood was collected for serum hormone measurements, and reproductive tissues were collected for histology and/or RNA extraction, as described below.

### Vaginal cytology

To determine the stages of the estrous cycle, vaginal cytology specimens were collected for 21–23 consecutive days, beginning at 10 weeks of age, using a cotton-tipped swab (Puritan Medical Product Company) wetted with normal saline solution (0.9% NaCl) and inserted into the vagina of the restrained mouse. The swab was gently turned and rolled against the vaginal wall and then removed. Cells were transferred to a dry glass slide by rolling the swab across the slide. The slide was air-dried and then stained with Three-Step Stain Kit (Richard-Allan Scientific) for 45 seconds. The slides were rinsed with distilled water, air-dried, and viewed under bright-field illumination at 20x magnification. The stages of the estrous cycle were determined based on the overall cellularity and the presence of nucleated epithelial cells (proestrus), cornified and anucleated epithelial cells (estrus), and/or leukocytes (diestrus), as described previously (Cora et al. 2015). One estrous cycle was defined as the sequential appearance of all stages, regardless of time spent in each. As it is not possible to know when the first stage began, estrous cycle lengths were calculated for each animal beginning with the appearance of the second stage and the percentage of time spent in each stage was determined for each completed cycle. For all genotypes, animals completed an average of ∼2 cycles during the assessment period.

### Hormone measurements

To determine circulating hormone levels, ∼750 μL of blood was collected by cardiac puncture of the ventricle while animals were under terminal anesthesia and serum LH and FSH concentrations were measured using immunofluorometric assays as described previously (Haavisto et al. 1993; Ahmed et al. 2017). Samples were collected from 27– 40-week-old female mice (n = 8 WT, 10 *Zeb1^200H^, and* 11 *Zeb1^200M^*) during proestrus, as determined by vaginal cytology.

### Histology

All tissues were collected from 27–40-week-old female mice during proestrus, as determined by vaginal cytology. Ovaries from WT*, Zeb1^200H^, and Zeb1^200M^* mice were fixed in 10% buffered formalin (Sigma) for 24 hours and stored in 70% Ethanol. Paraffin-embedded ovaries (one per animal, n = 8 WT, 10 *Zeb1^200H^, and* 11 *Zeb1^200M^* ovaries) were serially sectioned and stained with hematoxylin and eosin. For each section, two individuals blinded to genotype independently counted the number of early/primary follicles, defined as having less than two complete layers of granulosa cells, mature/secondary follicles, defined as having two or more complete layers of granulosa, and corpus luteum. These counts were summed for 2–5 sections taken from different levels of each ovary and then divided by the total area of those sections (mm^2^).

### RNA extraction

Total RNA was extracted from the pituitary, hypothalamus, and ovary tissues collected from 28–36-week-old WT, *Zeb1^200H^*, and *Zeb1^200M^* mice using TRI Reagent (ThermoFisher) according to the manufacturer’s protocol with the following modifications. Mouse tissues were rapidly dissected after euthanasia and flash frozen in Eppendorf tubes in liquid N_2_. Tissue was transferred to a 15 mL conical tube, 1–2 mL of TRI Reagent was added, and the tissue was homogenized with a TissueRuptor (Qiagen) and disposable probes. Samples were transferred to Eppendorf tubes (1 mL per tube) and phase separated with 200 μL chloroform (J.T. Baker Analytical). After isopropanol precipitation and two 70% ethanol washes, total RNA was resuspended in water.

### RNA-Seq and analysis

For all samples (n = 3–4 biological replicates per genotype), the TruSeq RNA Sample Prep Kit v2 (Illumina) was used to generate unstranded, poly(A)-selected RNA-seq libraries. Briefly, 1 μg of hypothalamus, pituitary, or ovary total RNA was poly(A) enriched and then reverse transcribed into double-strand cDNA. The cDNA samples were fragmented, end-repaired, and polyadenylated before ligation of TruSeq adapters containing an index sequence for multiplex sequencing. Multiplexing fragments containing TruSeq adapters on both ends were selectively enriched with 15 cycles of PCR. All libraries were sequenced on the Illumina HiSeq 2000 platform with 50 nt single-end reads.

Differential gene expression analysis of RNA-seq data was performed as follows. Reads were aligned to the mouse genome (mm10) using STAR v2.7.1a (Dobin et al. 2013) with the parameters “--outFilterType BySJout --outSAMtype BAM SortedByCoordinate -- readFilesCommand zcat --outFilterIntronMotifs RemoveNoncanonicalUnannotated”. For pituitary and hypothalamus libraries, aligned reads were assigned to genes using htseq-count v0.9.1 (Anders et al. 2015) with the parameters “-m union -s no” and annotations from Ensembl (Mus_musculus.GRCm38.87.gtf; downloaded May 15, 2017). For ovary libraries, aligned reads were assigned to genes using featureCounts (Liao et al. 2014) with the parameter “-s 0” and annotations from Gencode (m38.p6 gencode.vM25.basic.annotation.gtf; downloaded 7/10/24). Count files were merged to generate tables of counts for each tissue organized by genotype. Differential expression was determined using DESeq2 v1.18 (pituitary and hypothalamus) with the parameter “betaPrior = FALSE” and without the lfcShrink function or DESeq2 v1.38.3 (ovary) without the lfcShrink function (Love et al. 2014). RNA-seq browser tracks were visualized in the UCSC genome browser and IGV v2.3.34 (Robinson et al. 2011; Thorvaldsdottir et al. 2013). For counting pri-miR-200 species, reads mapping to *ENSMUSG00000086549* (miR-200b∼429 cluster) and *ENSMUSG00000087327* (miR-200c–141 cluster) were used.

For analysis of intron expression in the pituitary, STAR-aligned reads were assigned to gene introns using featureCounts (Liao et al. 2014) with the parameter “-s 0”, and annotations which were generated by loading the m38.p6 gencode.vM25.basic.annotation.gtf into the UCSC genome browser and downloading the corresponding intron structure information. For each gene, all introns were combined into a single annotation.

For RNA-seq plots (Fig. 3A–C, Supplemental Fig. S3B–G), only RNAs with a mean normalized CPM >1 in either WT or *Zeb1^200H^*samples (depending on the reference group for the comparison) are shown. Linear regression (Fig. 3D,I) was performed on genes with at least 50 raw read counts across samples.

Gene set enrichment analysis (GSEA) was performed using WebGestalt (Elizarraras et al. 2024) and gene ontology functional databases for biological processes and cell components with the following parameters: minimum number of IDs in category, 10; maximum number of IDs in category, 1000; significance level, FDR < 0.05; permutations, 1000. The input for GSEA was a list of M. musculus gene IDs with WT pituitary expression > 1 CPM ranked by STAT, which is the log2 fold change divided by log2 fold change standard error. Redundancy of genesets was reduced with the weighted set cover option.

Principal component analysis was performed on raw RNA sequencing read counts for all *Zeb1^200M^*, *Zeb1^200H^*, and WT samples of each tissue type. The Seurat functions NormalizeData, FindVariableFeatures (selecting the default of 2000 most variable genes), and ScaleData were implemented prior to principal component analysis (http://www.satijalab.org/seurat). The Seurat function RunPCA was used with the number of principal components equal to the total number of samples minus 1.

### Small-RNA sequencing and analysis

Small-RNA sequencing libraries were each prepared with 0.5 μg of total RNA from the pituitary of 28–36-week-old female mice (n = 3–4 samples per genotype). To minimize loss due to low starting input, 3 ug of total RNA from the budding yeast *Naumovozyma castelli* was added to each sample prior to size-selection. In addition, 0.5 fmoles of miR-427-5p (X*. tropicalis*) and 0.5 fmoles of miR-14 (*D. melanogaster*) were added as spike-ins. From each sample, RNAs that co-migrated within the range of 18 and 32 nt radiolabeled internal standards were isolated on a 15% polyacrylamide urea gel. Purified RNA was ligated to a preadenylated 3′ adapter (AppNNNNTCGTATGCCGTCTTCTGCTTGddC) using T4 RNA Ligase 2 KQ mutant (NEB) in a reaction supplemented with 10% polyethylene glycol (PEG 8000, NEB). To reduce ligation biases, this 3′ adapter had 4 random-sequence positions at its 5′ end. After gel purification on a 8% polyacrylamide urea gel, RNA was ligated to a 5′ adapter (GUUCAGAGUUCUACAGUCCGACGAUCNNNN) using T4 RNA Ligase I (NEB) in a reaction supplemented with 10% PEG. To reduce ligation biases, this adapter had 4 random-sequence positions at its 3′ end. After gel purification on an 8% polyacrylamide urea gel, RNA was reverse transcribed with SuperScript III (Invitrogen), and the cDNA was amplified ∼12 cycles with KAPA HiFi DNA polymerase (Kapa Biosystems). Amplified DNA was purified on a 3% metaphor agarose gel and submitted for sequencing. A step-by-step protocol for constructing libraries for small-RNA sequencing is available at http://bartellab.wi.mit.edu/protocols.html. Libraries were sequenced on the Illumina HiSeq platform with 50 nt single-end reads. Reads were trimmed of adaptor sequence using cutadapt (Martin 2011) and filtered using fastq_quality_filter (FastX Toolkit; http://hannonlab.cshl.edu/fastx_toolkit/) with the parameters “-q 30 –p 100” to ensure that all bases had an accuracy of 99.9%.

To count the miRNAs in each library, the first 18 nt of each read were string-matched to a dictionary of miRNA sequences. Mouse miRNA dictionaries were based on annotations in miRbase_v20. Matching to the first 18 nt has the advantage of combining all miRNA isoforms into a single count but also can result in ambiguous read assignments for mature miRNAs from distinct loci that are identical for the first 18 nt but differ at one or more positions closer to their 3′ ends (such as let-7a-5p and let-7c-5p). To avoid this ambiguity, these miRNAs (<5% of all annotations) were removed from the dictionary. Count files were merged to generate tables of counts organized by genotype. Differential expression was determined using DESeq2 (Love et al. 2014) with the parameter “betaPrior = FALSE”. For small-RNA sequencing plots (Fig. 4B; Supplemental Fig. S4A), only miRNAs with a mean normalized CPM >50 in WT samples and a CV <10 standard deviations above the median CV in both WT and mutant samples are shown. Similarly, linear regression (Supplemental Fig. S4B) was performed on miRNAs a mean normalized CPM >50 in WT samples.

### miRNA target analysis

miRNA target predictions were downloaded from TargetScanMouse Release 8.0 (McGeary et al. 2019). Differential expression data (DESeq2 output) was analyzed for repression of genes predicted to be miRNA targets. Only genes with at least 50 raw read counts across samples were considered. Three sets of predicted targets were analyzed: all predicted targets, conserved predicted targets, and top predicted targets. Top predicted targets were defined as the top 10% of all predicted targets based on cumulative weighted context++ scores (Agarwal et al. 2015). Each of the three sets of targets was compared to a control group of transcripts not predicted to be targets of the miRNA family in question. This nontarget set was selected as follows. First, 3′ UTR sequences from TargetScanMouse Release 8.0 (https://www.targetscan.org/cgi-bin/targetscan/data_download.cgi?db=mmu_72; downloaded on 7/10/24) were used to group genes into 10 bins based on 3′ UTR length. Next, for each target, one transcript not predicted to be a target of the miRNA family was selected, with replacement, from the corresponding UTR bin. For each target set, the distribution of log2 fold changes (computed using DESeq2) in *Zeb1^200M^* or *Zeb1^200H^* samples relative to WT samples was compared to the distribution of log2 fold changes of the 3′ UTR length-matched nontarget cohort, and statistical significance was determined using a Mann-Whitney test. The degree of repression is represented by subtracting the median log2 fold change of the target set from the median log2 fold change of its 3′ UTR length-matched nontarget set. The analysis described above was repeated 20 additional times for each target set, and the mean difference in median log2 fold changes and the median *p* value across the 21 replicates are reported (Fig. 4G; Supplemental Fig. S4E). For cumulative distribution functions displayed in Figures 4D, 4E and Supplemental Figure S4C and D, only the nontarget set corresponding to the all targets set is shown for simplicity.

### Quantification and statistical analysis

Graphs were generated in GraphPad Prism 7–10 or R and statistical analyses were performed using Excel, Graphpad, or R. Statistical parameters including the value of n, statistical test, and statistical significance (*p* value) are reported in the figures and their legends. For studies involving mouse tissues, replicates refer to samples derived from different mice. No statistical methods were used to predetermine sample size. Statistical tests were selected based on the desired comparison.

One-way ANOVA was used to assess significance when comparing fertility measurements between 3 genotypes; significant ANOVA results were followed by a post-hoc Kruskal-Wallis test. For differential expression of global measurements (RNA-seq, small-RNA sequencing), the DESeq2 software generated adjusted *p* values using the Wald test with the Benjamini-Hochberg procedure to correct for multiple-hypothesis testing. The Mann-Whitney test was used to compare cumulative distributions of mRNA fold changes between two gene sets. The Fisher’s exact test was used to determine the significance of overlap between gene sets.

### Data and software availability

Sequencing datasets generated in this study have been deposited in the GEO under accession numbers GSE279446 and GSE279447.

## ACKNOWLEDGMENTS

We thank A. Granger and members of the Bartel lab for helpful discussions; R. Bronson for assistance with histological analysis; J. Kero for performing the immunofluorometric assays; the Whitehead Institute Genome Technology Core for sequencing; the MIT histology core for tissue sectioning and staining; and the MIT Transgenic Facility for generating mutant mice. This research was supported by NIH grants R35GM118135 (D.P.B.), R35GM147463 (B.K.), T32GM144273 (L.E.E.), and T32GM136540 (L.E.E.) from the National Institute of General Medical Sciences and F30HL175923 (L.E.E.) from the National Heart, Lung, and Blood Institute. D.P.B is an investigator of the Howard Hughes Medical Institute.

## Author contributions

B.K., J.S, and D.P.B conceived the project and designed the study. S-J.H. generated the *Zeb1* and *Zeb2* knock-in mice. J.S. performed the breeding trial, collected tissue, prepared RNA-sequencing libraries, and analyzed histology. B.K. prepared small RNA sequencing libraries, analyzed histology, and analyzed sequencing data. L.E.E. analyzed sequencing data. B.K., J.S., and L.E.E. generated figures. B.K. drafted the manuscript and all authors revised the manuscript.

## REFERENCES

Aeschimann F, Neagu A, Rausch M, Großhans H. 2019. let-7 coordinates the transition to adulthood through a single primary and four secondary targets. Life Sci Alliance 2.

Agarwal V, Bell GW, Nam JW, Bartel DP. 2015. Predicting effective microRNA target sites in mammalian mRNAs. eLife 4.

Ahmed K, LaPierre MP, Gasser E, Denzler R, Yang Y, Rülicke T, Kero J, Latreille M, Stoffel M. 2017. Loss of microRNA-7a2 induces hypogonadotropic hypogonadism and infertility. The Journal of clinical investigation 127: 1061–1074.

Alon U. 2007. Network motifs: theory and experimental approaches. Nature reviews Genetics 8: 450–461.

Anders S, Pyl PT, Huber W. 2015. HTSeq--a Python framework to work with high-throughput sequencing data. *Bioinformatics (Oxford*, England) 31: 166–169.

Arnhold IJ, Lofrano-Porto A, Latronico AC. 2009. Inactivating mutations of luteinizing hormone beta-subunit or luteinizing hormone receptor cause oligo-amenorrhea and infertility in women. Horm Res 71: 75–82.

Balestrieri C, Alfarano G, Milan M, Tosi V, Prosperini E, Nicoli P, Palamidessi A, Scita G, Diaferia GR, Natoli G. 2018. Co-optation of Tandem DNA Repeats for the Maintenance of Mesenchymal Identity. Cell 173: 1150–1164.e1114.

Bartel DP. 2018. Metazoan MicroRNAs. Cell 173: 20–51.

Basciani S, Watanabe M, Mariani S, Passeri M, Persichetti A, Fiore D, Scotto d’Abusco A, Caprio M, Lenzi A, Fabbri A et al. 2012. Hypogonadism in a patient with two novel mutations of the luteinizing hormone β-subunit gene expressed in a compound heterozygous form. J Clin Endocrinol Metab 97: 3031–3038.

Bracken CP, Goodall GJ, Gregory PA. 2024. RNA regulatory mechanisms controlling TGF-β signaling and EMT in cancer. Seminars in Cancer Biology 102-103: 4–16.

Bracken CP, Gregory PA, Kolesnikoff N, Bert AG, Wang J, Shannon MF, Goodall GJ. 2008. A double-negative feedback loop between ZEB1-SIP1 and the microRNA-200 family regulates epithelial-mesenchymal transition. Cancer research 68: 7846–7854.

Burk U, Schubert J, Wellner U, Schmalhofer O, Vincan E, Spaderna S, Brabletz T. 2008. A reciprocal repression between ZEB1 and members of the miR-200 family promotes EMT and invasion in cancer cells. EMBO reports 9: 582–589.

Carson SA, Kallen AN. 2021. Diagnosis and Management of Infertility: A Review. Jama 326: 65–76.

Cassidy JJ, Jha AR, Posadas DM, Giri R, Venken KJ, Ji J, Jiang H, Bellen HJ, White KP, Carthew RW. 2013. miR-9a minimizes the phenotypic impact of genomic diversity by buffering a transcription factor. Cell 155: 1556–1567.

Chua HL, Bhat-Nakshatri P, Clare SE, Morimiya A, Badve S, Nakshatri H. 2007. NF-κB represses E-cadherin expression and enhances epithelial to mesenchymal transition of mammary epithelial cells: potential involvement of ZEB-1 and ZEB-2. Oncogene 26: 711–724.

Cora MC, Kooistra L, Travlos G. 2015. Vaginal Cytology of the Laboratory Rat and Mouse: Review and Criteria for the Staging of the Estrous Cycle Using Stained Vaginal Smears. Toxicol Pathol 43: 776–793.

Dalkin AC, Haisenleder DJ, Ortolano GA, Ellis TR, Marshall JC. 1989. The Frequency of Gonadotropin-Releasing-Hormone Stimulation Differentially Regulates Gonadotropin Subunit Messenger Ribonucleic Acid Expression*. Endocrinology 125: 917–923.

Dobin A, Davis CA, Schlesinger F, Drenkow J, Zaleski C, Jha S, Batut P, Chaisson M, Gingeras TR. 2013. STAR: ultrafast universal RNA-seq aligner. *Bioinformatics (Oxford*, England) 29: 15–21.

Dorsett Y, McBride KM, Jankovic M, Gazumyan A, Thai TH, Robbiani DF, Di Virgilio M, Reina San-Martin B, Heidkamp G, Schwickert TA et al. 2008. MicroRNA-155 suppresses activation-induced cytidine deaminase-mediated Myc-Igh translocation. Immunity 28: 630–638.

Drexel T, Mahofsky K, Latham R, Zimmer M, Cochella L. 2016. Neuron type-specific miRNA represses two broadly expressed genes to modulate an avoidance behavior in C. elegans. Genes & development 30: 2042–2047.

Ecsedi M, Rausch M, Großhans H. 2015. The let-7 microRNA directs vulval development through a single target. Developmental cell 32: 335–344.

Elizarraras JM, Liao Y, Shi Z, Zhu Q, Pico AR, Zhang B. 2024. WebGestalt 2024: faster gene set analysis and new support for metabolomics and multi-omics. Nucleic acids research 52: W415–w421.

Ferrell JE, Jr., Ha SH. 2014. Ultrasensitivity part III: cascades, bistable switches, and oscillators. Trends Biochem Sci 39: 612–618.

Friedman RC, Farh KK, Burge CB, Bartel DP. 2009. Most mammalian mRNAs are conserved targets of microRNAs. Genome research 19: 92–105.

Garaulet DL, Zhang B, Wei L, Li E, Lai EC. 2020. miRNAs and Neural Alternative Polyadenylation Specify the Virgin Behavioral State. Developmental cell 54: 410–423.e414.

Gregory PA, Bert AG, Paterson EL, Barry SC, Tsykin A, Farshid G, Vadas MA, Khew-Goodall Y, Goodall GJ. 2008. The miR-200 family and miR-205 regulate epithelial to mesenchymal transition by targeting ZEB1 and SIP1. Nature cell biology 10: 593–601.

Gregory PA, Bracken CP, Smith E, Bert AG, Wright JA, Roslan S, Morris M, Wyatt L, Farshid G, Lim YY et al. 2011. An autocrine TGF-beta/ZEB/miR-200 signaling network regulates establishment and maintenance of epithelial-mesenchymal transition. Mol Biol Cell 22: 1686–1698.

Gröger CJ, Grubinger M, Waldhör T, Vierlinger K, Mikulits W. 2012. Meta-analysis of gene expression signatures defining the epithelial to mesenchymal transition during cancer progression. PloS one 7: e51136.

Guan T, Dominguez CX, Amezquita RA, Laidlaw BJ, Cheng J, Henao-Mejia J, Williams A, Flavell RA, Lu J, Kaech SM. 2018. ZEB1, ZEB2, and the miR-200 family form a counterregulatory network to regulate CD8(+) T cell fates. J Exp Med 215: 1153–1168.

Haavisto AM, Pettersson K, Bergendahl M, Perheentupa A, Roser JF, Huhtaniemi I. 1993. A supersensitive immunofluorometric assay for rat luteinizing hormone. Endocrinology 132: 1687–1691.

Haisenleder DJ, Dalkin AC, Ortolano GA, Marshall JC, Shupnik MA. 1991. A Pulsatile Gonadotropin-Releasing Hormone Stimulus is Required to Increase Transcription of the Gonadotropin Subunit Genes: Evidence for Differential Regulation of Transcription by Pulse Frequency In Vivo*. Endocrinology 128: 509–517.

Halvorson LM, Ito M, Jameson JL, Chin WW. 1998. Steroidogenic factor-1 and early growth response protein 1 act through two composite DNA binding sites to regulate luteinizing hormone beta-subunit gene expression. The Journal of biological chemistry 273: 14712–14720.

Halvorson LM, Kaiser UB, Chin WW. 1996. Stimulation of luteinizing hormone beta gene promoter activity by the orphan nuclear receptor, steroidogenic factor-1. The Journal of biological chemistry 271: 6645–6650.

Hannon CE, Blythe SA, Wieschaus EF. 2017. Concentration dependent chromatin states induced by the bicoid morphogen gradient. eLife 6.

Hasuwa H, Ueda J, Ikawa M, Okabe M. 2013. miR-200b and miR-429 function in mouse ovulation and are essential for female fertility. Science 341: 71–73.

Heinz S, Benner C, Spann N, Bertolino E, Lin YC, Laslo P, Cheng JX, Murre C, Singh H, Glass CK. 2010. Simple combinations of lineage-determining transcription factors prime cis-regulatory elements required for macrophage and B cell identities. Molecular cell 38: 576–589.

Hurtado A, Mota-Gómez I, Lao M, Real FM, Jedamzick J, Burgos M, Lupiáñez DG, Jiménez R, Barrionuevo FJ. 2024. Complete male-to-female sex reversal in XY mice lacking the miR-17∼92 cluster. Nature communications 15: 3809.

Hurteau GJ, Carlson JA, Spivack SD, Brock GJ. 2007. Overexpression of the microRNA hsa-miR-200c leads to reduced expression of transcription factor 8 and increased expression of E-cadherin. Cancer research 67: 7972–7976.

Janati-Idrissi S, de Abreu MR, Guyomar C, de Mello F, Nguyen T, Mechkouri N, Gay S, Montfort J, Gonzalez AA, Abbasi M et al. 2024. Looking for a needle in a haystack: de novo phenotypic target identification reveals Hippo pathway-mediated miR-202 regulation of egg production. Nucleic acids research 52: 738–754.

Jin L, Liu Y, Wu Y, Huang Y, Zhang D. 2023. REST Is Not Resting: REST/NRSF in Health and Disease. Biomolecules 13.

Kahlert UD, Maciaczyk D, Doostkam S, Orr BA, Simons B, Bogiel T, Reithmeier T, Prinz M, Schubert J, Niedermann G et al. 2012. Activation of canonical WNT/β-catenin signaling enhances in vitro motility of glioblastoma cells by activation of ZEB1 and other activators of epithelial-to-mesenchymal transition. Cancer letters 325: 42–53.

Kaiser UB, Jakubowiak A, Steinberger A, Chin WW. 1997. Differential effects of gonadotropin-releasing hormone (GnRH) pulse frequency on gonadotropin subunit and GnRH receptor messenger ribonucleic acid levels in vitro. Endocrinology 138: 1224–1231.

Keenan AB, Torre D, Lachmann A, Leong AK, Wojciechowicz ML, Utti V, Jagodnik KM, Kropiwnicki E, Wang Z, Ma’ayan A. 2019. ChEA3: transcription factor enrichment analysis by orthogonal omics integration. Nucleic acids research 47: W212–w224.

Korpal M, Kang Y. 2008. The emerging role of miR-200 family of MicroRNAs in epithelial-mesenchymal transition and cancer metastasis. RNA biology 5: 115–119.

Korpal M, Lee ES, Hu G, Kang Y. 2008. The miR-200 family inhibits epithelial-mesenchymal transition and cancer cell migration by direct targeting of E-cadherin transcriptional repressors ZEB1 and ZEB2. The Journal of biological chemistry 283: 14910–14914.

Kuzniewska B, Rejmak K, Nowacka A, Ziółkowska M, Milek J, Magnowska M, Gruchota J, Gewartowska O, Borsuk E, Salamian A et al. 2022. Disrupting interaction between miR-132 and Mmp9 3’UTR improves synaptic plasticity and memory in mice. Frontiers in molecular neuroscience 15: 924534.

Lai EC, Posakony JW. 1997. The Bearded box, a novel 3’ UTR sequence motif, mediates negative post-transcriptional regulation of Bearded and Enhancer of split Complex gene expression. Development (Cambridge, England) 124: 4847–4856.

Liao Y, Smyth GK, Shi W. 2014. featureCounts: an efficient general purpose program for assigning sequence reads to genomic features. Bioinformatics (Oxford, England) 30: 923–930.

Lofrano-Porto A, Barra GB, Giacomini LA, Nascimento PP, Latronico AC, Casulari LA, da Rocha Neves Fde A. 2007. Luteinizing hormone beta mutation and hypogonadism in men and women. N Engl J Med 357: 897–904.

Love MI, Huber W, Anders S. 2014. Moderated estimation of fold change and dispersion for RNA-seq data with DESeq2. Genome biology 15: 550.

Lu LF, Gasteiger G, Yu IS, Chaudhry A, Hsin JP, Lu Y, Bos PD, Lin LL, Zawislak CL, Cho S et al. 2015. A Single miRNA-mRNA Interaction Affects the Immune Response in a Context- and Cell-Type-Specific Manner. Immunity 43: 52–64.

Lunenfeld B. 2012. Gonadotropin stimulation: past, present and future. Reprod Med Biol 11: 11–25.

Ma X, Dong Y, Matzuk MM, Kumar TR. 2004. Targeted disruption of luteinizing hormone beta-subunit leads to hypogonadism, defects in gonadal steroidogenesis, and infertility. Proceedings of the National Academy of Sciences of the United States of America 101: 17294–17299.

Martin M. 2011. Cutadapt removes adapter sequences from high-throughput sequencing reads. 2011 17.

McGeary SE, Lin KS, Shi CY, Pham TM, Bisaria N, Kelley GM, Bartel DP. 2019. The biochemical basis of microRNA targeting efficacy. Science 366.

McJunkin K, Ambros V. 2017. A microRNA family exerts maternal control on sex determination in C. elegans. Genes & development 31: 422–437.

Messina A, Langlet F, Chachlaki K, Roa J, Rasika S, Jouy N, Gallet S, Gaytan F, Parkash J, Tena-Sempere M et al. 2016. A microRNA switch regulates the rise in hypothalamic GnRH production before puberty. Nature neuroscience 19: 835–844.

Mikhael S, Punjala-Patel A, Gavrilova-Jordan L. 2019. Hypothalamic-Pituitary-Ovarian Axis Disorders Impacting Female Fertility. Biomedicines 7.

Mildner A, Chapnik E, Varol D, Aychek T, Lampl N, Rivkin N, Bringmann A, Paul F, Boura-Halfon S, Hayoun YS et al. 2017. MicroRNA-142 controls thymocyte proliferation. Eur J Immunol 47: 1142–1152.

Munro MG, Balen AH, Cho S, Critchley HOD, Díaz I, Ferriani R, Henry L, Mocanu E, van der Spuy ZM, Disorders ftFCoM, et al. 2022. The FIGO ovulatory disorders classification system. International Journal of Gynecology & Obstetrics 159: 1–20.

Naqvi S, Kim S, Hoskens H, Matthews HS, Spritz RA, Klein OD, Hallgrímsson B, Swigut T, Claes P, Pritchard JK et al. 2023. Precise modulation of transcription factor levels identifies features underlying dosage sensitivity. Nature genetics 55: 841–851.

Park SM, Gaur AB, Lengyel E, Peter ME. 2008. The miR-200 family determines the epithelial phenotype of cancer cells by targeting the E-cadherin repressors ZEB1 and ZEB2. Genes & development 22: 894–907.

Pierce JG, Parsons TF. 1981. Glycoprotein hormones: structure and function. Annu Rev Biochem 50: 465–495.

Renthal NE, Chen CC, Williams KC, Gerard RD, Prange-Kiel J, Mendelson CR. 2010. miR-200 family and targets, ZEB1 and ZEB2, modulate uterine quiescence and contractility during pregnancy and labor. Proceedings of the National Academy of Sciences of the United States of America 107: 20828–20833.

Risma KA, Clay CM, Nett TM, Wagner T, Yun J, Nilson JH. 1995. Targeted overexpression of luteinizing hormone in transgenic mice leads to infertility, polycystic ovaries, and ovarian tumors. Proceedings of the National Academy of Sciences of the United States of America 92: 1322–1326.

Robinson JT, Thorvaldsdottir H, Winckler W, Guttman M, Lander ES, Getz G, Mesirov JP. 2011. Integrative genomics viewer. Nature biotechnology 29: 24–26.

Ruf-Zamojski F, Zhang Z, Zamojski M, Smith GR, Mendelev N, Liu H, Nudelman G, Moriwaki M, Pincas H, Castanon RG et al. 2021. Single nucleus multi-omics regulatory landscape of the murine pituitary. Nature communications 12: 2677.

Saitoh M. 2023. Transcriptional regulation of EMT transcription factors in cancer. Semin Cancer Biol 97: 21–29.

Schultz H, Zhou X, Alonso CAI, Ongaro L, Lin Y-F, Loka M, Brabletz T, Brabletz S, Stemmler MP, Boehm U et al. 2024. ZEB1 Inhibits LHβ Subunit Transcription When Overexpressed, but Is Dispensable for LH Synthesis in Mice. Endocrinology 165.

Song JW, Hwang HJ, Lee CM, Park GH, Kim CS, Lee SJ, Ihm SH. 2018. Hypogonadotrophic hypogonadism due to a mutation in the luteinizing hormone β-subunit gene. Korean J Intern Med 33: 638–641.

Szabó F, Köves K, Gál L. 2024. History of the Development of Knowledge about the Neuroendocrine Control of Ovulation—Recent Knowledge on the Molecular Background. in International journal of molecular sciences.

Teng G, Hakimpour P, Landgraf P, Rice A, Tuschl T, Casellas R, Papavasiliou FN. 2008. MicroRNA-155 is a negative regulator of activation-induced cytidine deaminase. Immunity 28: 621–629.

Thompson IR, Kaiser UB. 2014. GnRH pulse frequency-dependent differential regulation of LH and FSH gene expression. Mol Cell Endocrinol 385: 28–35.

Thorvaldsdottir H, Robinson JT, Mesirov JP. 2013. Integrative Genomics Viewer (IGV): high-performance genomics data visualization and exploration. Briefings in bioinformatics 14: 178–192.

Title AC, Hong SJ, Pires ND, Hasenöhrl L, Godbersen S, Stokar-Regenscheit N, Bartel DP, Stoffel M. 2018. Genetic dissection of the miR-200-Zeb1 axis reveals its importance in tumor differentiation and invasion. Nature communications 9: 4671.

Title AC, Silva PN, Godbersen S, Hasenöhrl L, Stoffel M. 2021. The miR-200-Zeb1 axis regulates key aspects of β-cell function and survival in vivo. Mol Metab 53: 101267.

Tremblay JJ, Drouin J. 1999. Egr-1 is a downstream effector of GnRH and synergizes by direct interaction with Ptx1 and SF-1 to enhance luteinizing hormone beta gene transcription. Mol Cell Biol 19: 2567–2576.

Truett GE, Heeger P, Mynatt RL, Truett AA, Walker JA, Warman ML. 2000. Preparation of PCR-quality mouse genomic DNA with hot sodium hydroxide and tris (HotSHOT). BioTechniques 29: 52, 54.

Valdes-Socin H, Salvi R, Daly AF, Gaillard RC, Quatresooz P, Tebeu PM, Pralong FP, Beckers A. 2004. Hypogonadism in a patient with a mutation in the luteinizing hormone beta-subunit gene. N Engl J Med 351: 2619–2625.

Vannier C, Mock K, Brabletz T, Driever W. 2013. Zeb1 Regulates E-cadherin and Epcam (Epithelial Cell Adhesion Molecule) Expression to Control Cell Behavior in Early Zebrafish Development*. Journal of Biological Chemistry 288: 18643–18659.

Weiss J, Axelrod L, Whitcomb RW, Harris PE, Crowley WF, Jameson JL. 1992. Hypogonadism caused by a single amino acid substitution in the beta subunit of luteinizing hormone. N Engl J Med 326: 179–183.

Wolfe MW, Call GB. 1999. Early growth response protein 1 binds to the luteinizing hormone-beta promoter and mediates gonadotropin-releasing hormone-stimulated gene expression. Mol Endocrinol 13: 752–763.

Yang B, McJunkin K. 2020. The mir-35-42 binding site in the nhl-2 3’UTR is dispensable for development and fecundity. MicroPubl Biol 2020.

Yang B, Schwartz M, McJunkin K. 2020. In vivo CRISPR screening for phenotypic targets of the mir-35-42 family in C. elegans. Genes & development 34: 1227–1238.

Yang X, Ochin H, Shu L, Liu J, Shen J, Liu J, Lin C, Cui Y. 2018. Homozygous nonsense mutation Trp28X in the LHB gene causes male hypogonadism. J Assist Reprod Genet 35: 913–919.

Young C, Caffrey M, Janton C, Kobayashi T. 2022. Reversing the miRNA -5p/-3p stoichiometry reveals physiological roles and targets of miR-140 miRNAs. Rna 28: 854–864.

Zhang L, Zhang W, Li Y, Alvarez A, Li Z, Wang Y, Song L, Lv D, Nakano I, Hu B et al. 2016. SHP-2-upregulated ZEB1 is important for PDGFRα-driven glioma epithelial– mesenchymal transition and invasion in mice and humans. Oncogene 35: 5641–5652.

