## Supplemental Table 1 and Figures 1-5 for "Derepression of a single microRNA target causes female infertility in mice"

| **Offspring of *Zeb1 6B6*$^{200H}$ x *Zeb1 6B6*$^{200H}$ Cross** | | | |
|---|---|---|---|
| | *Zeb1 6B6*$^{+/+}$ | *Zeb1 6B6*$^{200H}$ | *Zeb1 6B6*$^{200M}$ |
| Observed (P21) | 36 | 62 | 20 |
| % observed | 31 | 53 | 17 |
| % expected | 25 | 50 | 25 |

Shown are results of crossing heterozygous mice on a C57Bl/6J background (>15 backcrosses) derived from targeted ESC clone 6B6. Statistical significance was determined by $X^2$ test (p = 0.10).

| **Offspring of *Zeb1 3C5*$^{200H}$ x *Zeb1 3C5*$^{200H}$ Cross** | | | |
|---|---|---|---|
| | *Zeb1 3C5*$^{+/+}$ | *Zeb1 3C5*$^{200H}$ | *Zeb1 3C5*$^{200M}$ |
| Observed (P21) | 25 | 43 | 16 |
| % observed | 30 | 51 | 19 |
| % expected | 25 | 50 | 25 |

Shown are results of crossing heterozygous mice on a C57Bl/6J background (>15 backcrosses) derived from targeted ESC clone 3C5. Statistical significance was determined by $X^2$ test (p = 0.37).

**Supplemental Table S1. Viability of mutant mice generated in this study, related to Figure 1.**

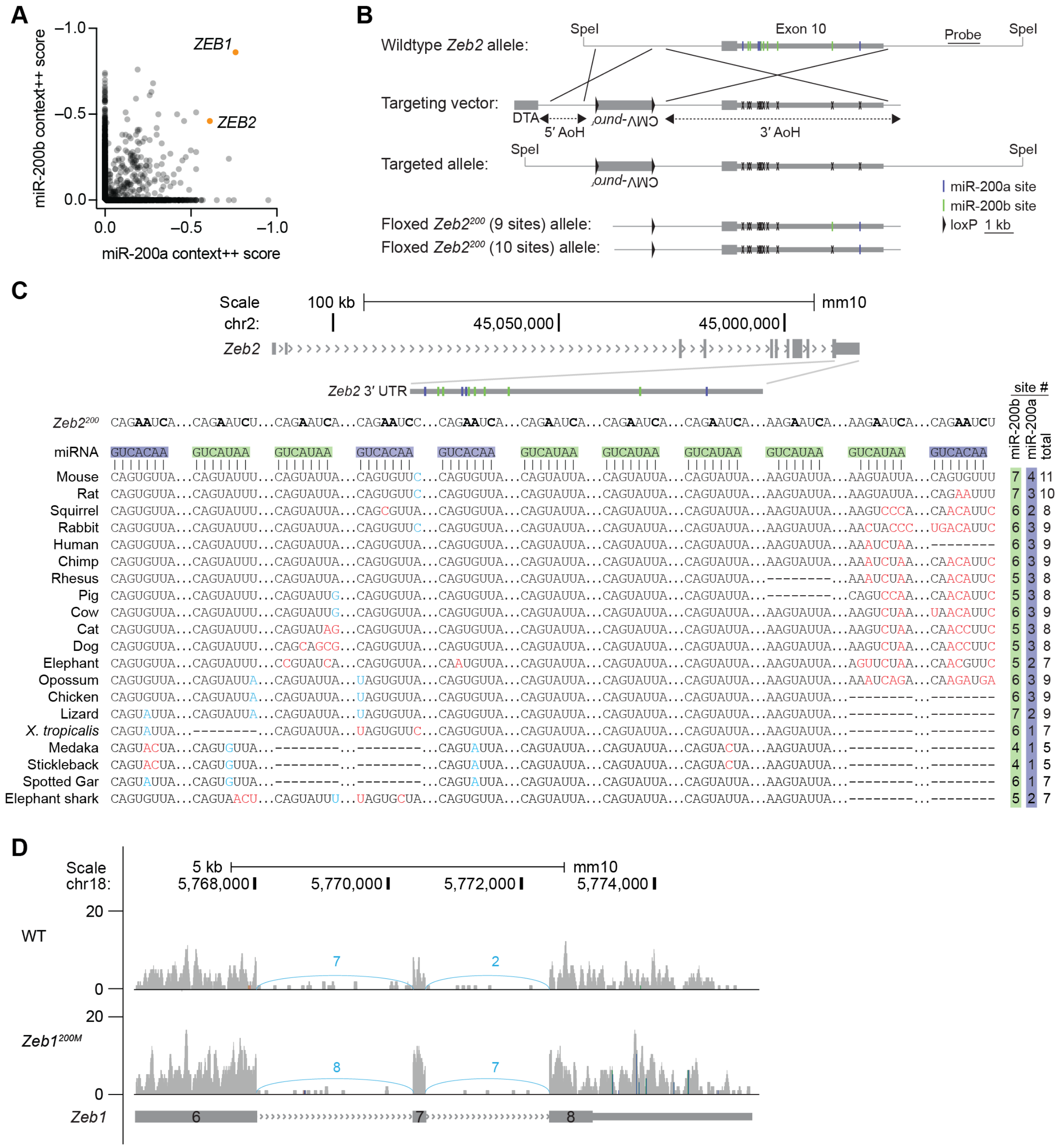


**Supplemental Figure S1. *Zeb1* is a top target of the miR-200 superfamily, related to Figure 1.** (A) Predicted targets of miR-200b/c/429 and miR-141/200a in humans. Otherwise, as in Figure 1B. (B) *Zeb2* gene targeting strategy. Shown between the wild-type exon 10 and the recombinant alleles is the vector used to insert 2–3 point mutations into each miR-200a/b site (gray boxes, exons; green lines, miR-200b sites; blue lines, miR-200a sites; X, mutant sites; filled triangles, loxP sites; CMV-*puro*[r], puromycin-resistance cassette; DTA, diphtheria toxin A; AoH, arm of homology). Screening of ES cell clones was performed by long-range PCR as well as Southern blot of SpeI-digested genomic DNA with a 3′ probe downstream of exon 10. (C) Organization of the murine *Zeb2* locus. The *Zeb2* gene model (gray boxes, exons; > > >, introns) is depicted with an inset of the 3′ UTR indicating the location of the binding sites for miR-200b (green boxes) and miR-200a (blue boxes). Diagramed below that is the pairing of seven miR-200b sites and four miR-200a sites as well the mutations that were introduced in each binding site to disrupt miRNA binding. Otherwise, as in Figure 1C. (D) The minimal influence of miRNA binding site mutations in the *Zeb1* 3′ UTR on splicing and 3′ end formation. RNA-seq coverage tracks for wild-type and *Zeb1*$^{200M}$ pituitary (*y*-axis, average number of reads per genomic window) derived from libraries with similar sequencing depths (~11 million reads for each library) and visualized in IGV are shown above the last three exons of the *Zeb1* gene model (gray boxes, exons; > > >, introns). Single nucleotide mismatches to the reference sequence with an allele frequency of at least 50% are indicated in blue/green and reflect successful mutagenesis of miR-200a/b binding sites. Splice junctions are represented by arcs with the numbers above each arc refer to unique splice junction mapping reads.

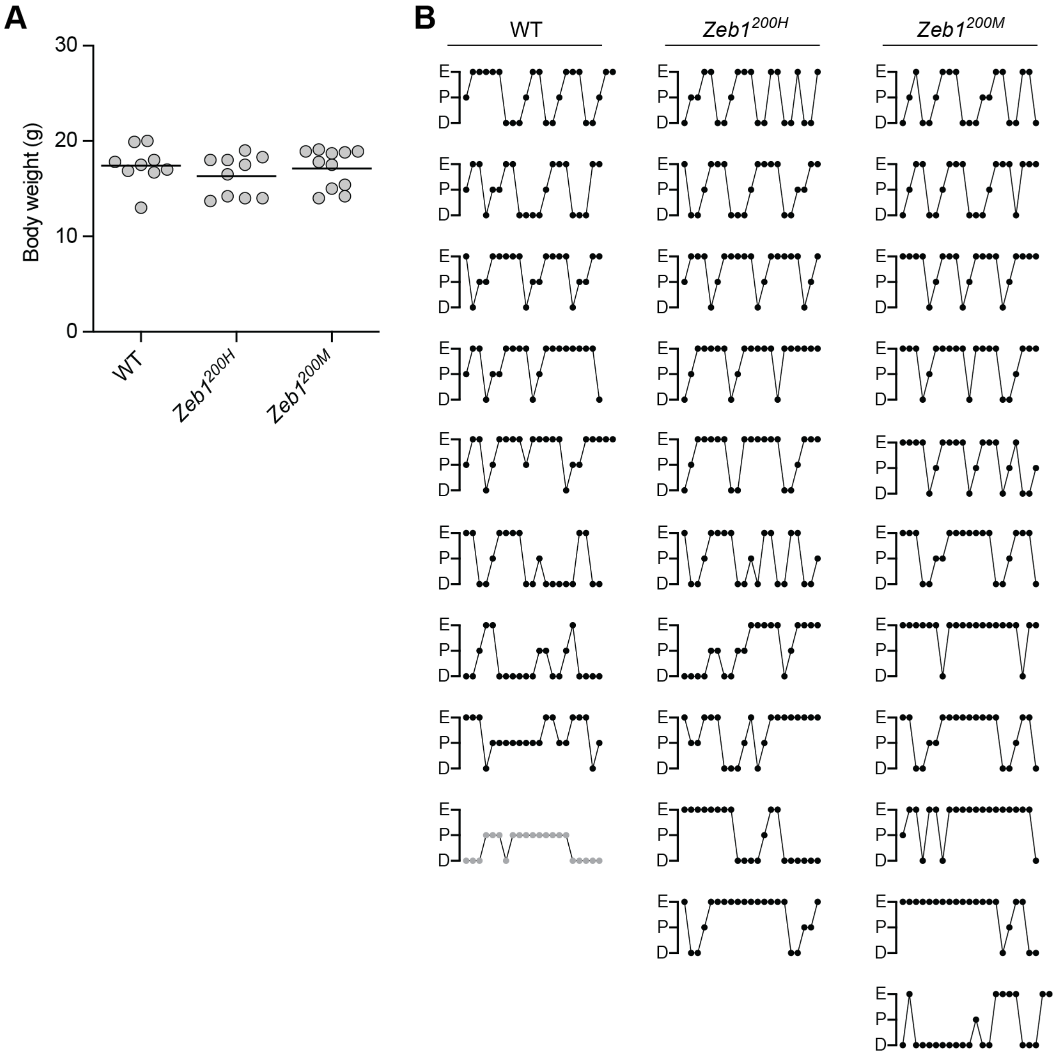


**Supplemental Figure S2. Female *Zeb1*$^{200M}$ mice have decreased fertility, related to Figure 2.** (A) Influence of *Zeb1*$^{200}$ allele on body weight. Plotted are weights (in grams) for WT, *Zeb1*$^{200H}$, and *Zeb1*$^{200M}$ female mice at 10 weeks of age (black line, mean; n = 9–11 mice per genotype). No statistically significant changes (adjusted $p$ value < 0.05) were detected (ANOVA with Kruskal-Wallis test). (B) Influence of *Zeb1*$^{200}$ alleles on estrus cycle length and stage. Plotted are consecutive measurements of estrus cycle stage, as determined by vaginal cytology, beginning at 10 weeks of age for 9 WT, 10 *Zeb1*$^{200H}$, and 11 *Zeb1*$^{200M}$ female mice. Each dot represents a day (n = 21–23 days per animal; E, estrus; P, proestrus; D, diestrus). Of the 30 mice in our cohort, 29 completed at least one estrus cycle and were included in the subsequent breeding trial. One WT mouse (gray dots) never cycled and was excluded from further studies.

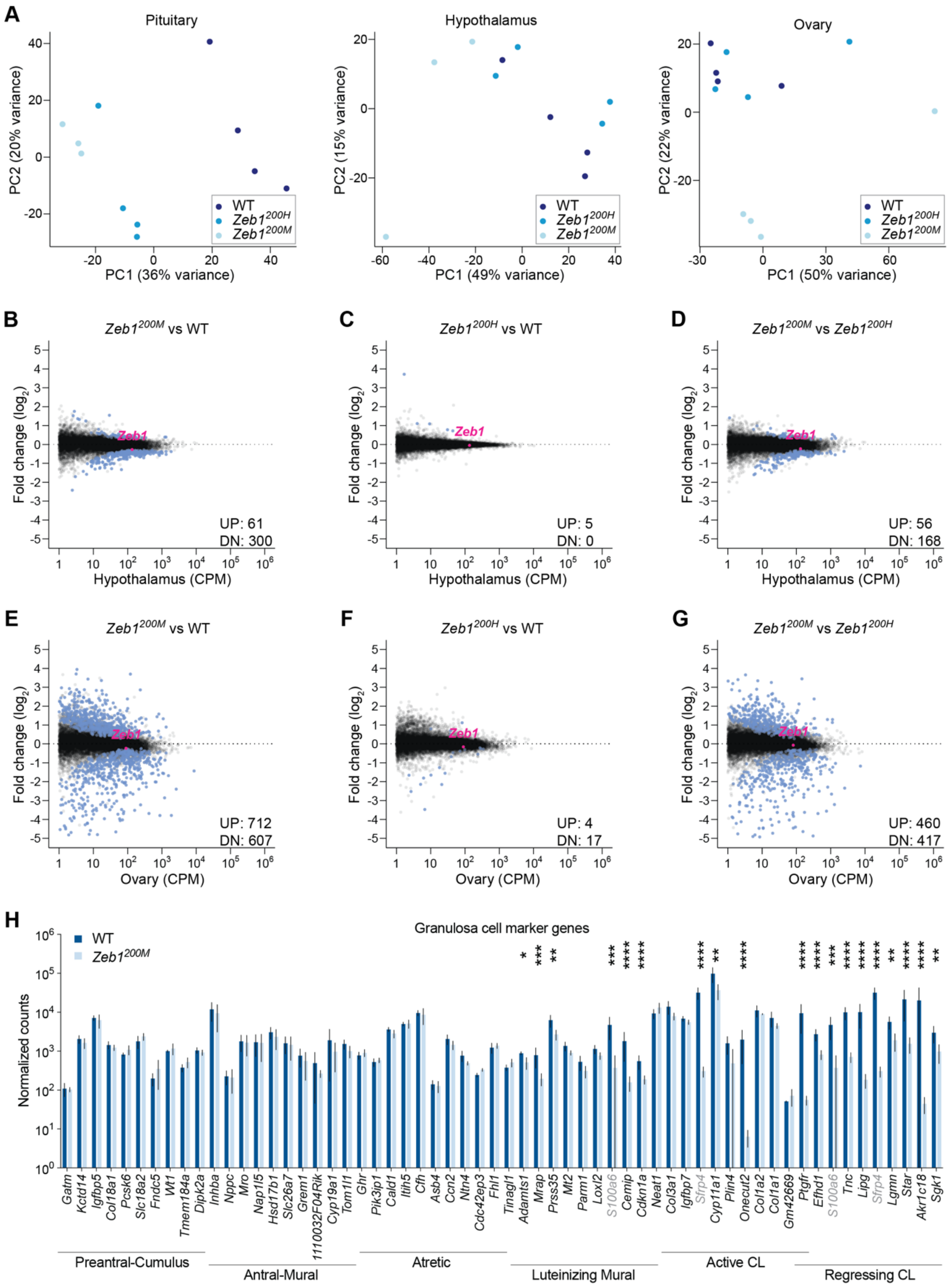
A
Pituitary
Hypothalamus
Ovary
PC2 (20% variance)
PC2 (15% variance)
PC2 (22% variance)
PC1 (36% variance)
PC1 (49% variance)
PC1 (50% variance)
WT
Zeb1^200H
Zeb1^200M
B
Zeb1^200M vs WT
C
Zeb1^200H vs WT
D
Zeb1^200M vs Zeb1^200H
Fold change (log2)
Zeb1
UP: 61
DN: 300
UP: 5
DN: 0
UP: 56
DN: 168
Hypothalamus (CPM)
E
Zeb1^200M vs WT
F
Zeb1^200H vs WT
G
Zeb1^200M vs Zeb1^200H
UP: 712
DN: 607
UP: 4
DN: 17
UP: 460
DN: 417
Ovary (CPM)
H
Granulosa cell marker genes
WT
Zeb1^200M
Normalized counts
Gatm Kctd14 Igfbp5 Col18a1 Pcsk6 Slc18a2 Fndc5 Wt1 Tmem184a Dipk2a
Inhba Nppc Mro Nap1l5 Hsd17b1 Slc26a7 Grem1 1110032F04Rik Cyp19a1 Tom1l1
Ghr Pik3ip1 Cald1 Itih5 Cfh Asb4 Ccn2 Ntn4 Cdc42ep3 Fhl1
Tinagl1 Adamts1 Mrap Prss35 Mt2 Parm1 Loxl2 S100a6 Cemip Cdkn1a
Neat1 Col3a1 Igfbp7 Sfrp4 Cyp11a1 Plin4 Onecut2 Col1a2 Col1a1 Gm42669
Ptgfr Efhd1 S100a6 Tnc Lipg Sfrp4 Lgmn Star Akr1c18 Sgk1
Preantral-Cumulus
Antral-Mural
Atretic
Luteinizing Mural
Active CL
Regressing CL

**Supplemental Figure S3. Widespread gene expression changes induced by modest derepression of *Zeb1*, related to Figure 3.** (A) Principal component analysis of WT and *Zeb1*$^{200}$ mutant tissues. Plotted are the first two principal components (PC1, PC2) for pituitary (left), hypothalamus (center), and ovary (right) based on analysis of the 2000 most variable genes derived from RNA sequencing libraries. Each circle represents a unique sample/library and each genotype is represented by a different color (WT, dark blue; *Zeb1*$^{200H}$, blue; *Zeb1*$^{200M}$, light blue). (B–D) The influence of *Zeb1*$^{200}$ alleles on RNA levels in the hypothalamus. Otherwise, as in Figure 3A–C. (E–G) The influence of *Zeb1*$^{200}$ alleles on RNA levels in the ovary. Otherwise, as in Figure 3A–C. (H) The influence of *Zeb1*$^{200M}$ on expression of granulosa cell marker genes in the ovary. Plotted are the mean counts per million mapped reads (CPM), as determined by RNA sequencing, for 60 genes previously attributed to 6 different granulosa cell types (Morris et al. 2022) in WT and *Zeb1*$^{200M}$ ovary (WT, dark blue; *Zeb1*$^{200M}$, light blue; CL, corpus lutea; error bars, standard deviation; n = 3–4 per genotype). Benjamini-Hochberg adjusted *p* values, as determined by a Wald test (DESeq2), are indicated (*, $p < 0.05$; **, $p < 0.01$, ***, $p < 0.001$, ****, $p < 0.0001$). Two genes, *S100a6* and *Sfrp4*, indicated by gray text are attributed to two different granulosa cell types.

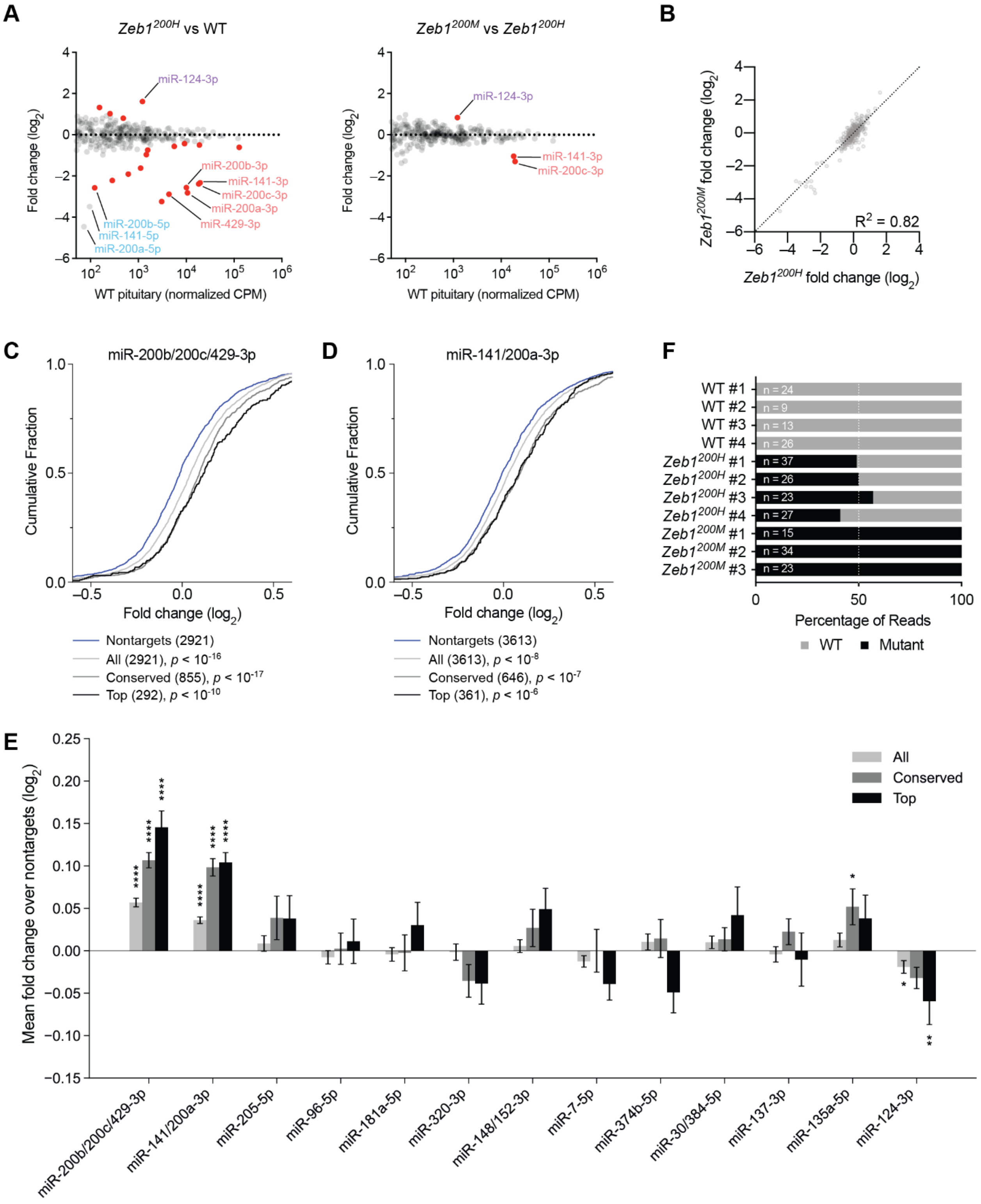
A
Zeb1200H vs WT
Zeb1200M vs Zeb1200H
Fold change (log2)
WT pituitary (normalized CPM)
miR-124-3p
miR-200b-3p
miR-141-3p
miR-200c-3p
miR-200a-3p
miR-429-3p
miR-200b-5p
miR-141-5p
miR-200a-5p
B
Zeb1200M fold change (log2)
Zeb1200H fold change (log2)
R2 = 0.82
C
miR-200b/200c/429-3p
Cumulative Fraction
Fold change (log2)
Nontargets (2921)
All (2921), p < 10-16
Conserved (855), p < 10-17
Top (292), p < 10-10
D
miR-141/200a-3p
Cumulative Fraction
Fold change (log2)
Nontargets (3613)
All (3613), p < 10-8
Conserved (646), p < 10-7
Top (361), p < 10-6
F
WT #1
WT #2
WT #3
WT #4
Zeb1200H #1
Zeb1200H #2
Zeb1200H #3
Zeb1200H #4
Zeb1200M #1
Zeb1200M #2
Zeb1200M #3
n = 24
n = 9
n = 13
n = 26
n = 37
n = 26
n = 23
n = 27
n = 15
n = 34
n = 23
Percentage of Reads
WT
Mutant
E
Mean fold change over nontargets (log2)
All
Conserved
Top
miR-200b/200c/429-3p
miR-141/200a-3p
miR-205-5p
miR-96-5p
miR-181a-5p
miR-320-3p
miR-148/152-3p
miR-7-5p
miR-374b-5p
miR-30/384-5p
miR-137-3p
miR-135a-5p
miR-124-3p

**Supplemental Figure S4. Increased Zeb1 causes decreased expression of miR-200a/b and increased expression of miR-200a/b targets, related to Figure 4.** (A) The influence of *Zeb1*$^{200}$ alleles on mature miRNA levels in the pituitary. Plotted are fold changes in mean miRNA levels for *Zeb1*$^{200H}$ pituitary relative to wild-type pituitary (left) and *Zeb1*$^{200M}$ pituitary relative to *Zeb1*$^{200H}$ pituitary (right), as determined by small-RNA sequencing and plotted as a function of expression in wild-type pituitary (n = 3–4 per genotype). Otherwise, as in Figure 4B. (B) Correlation of miRNA fold changes between *Zeb1*$^{200}$ alleles. Otherwise, as in Figure 3D. (C–D) The influence of *Zeb1*$^{200H}$ relative to wild-type on expression of miR-200b/200c/429-3p and miR-141/200a-3p predicted targets. Otherwise, as in Figure 4D–E. (E) The influence of *Zeb1*$^{200H}$ on expression of predicted targets of differentially expressed miRNAs. Otherwise, as in Figure 4F. (F) Allele-specific expression of *Zeb1* in wild-type and mutant pituitary. Plotted are the percentage of reads matching the wild-type or mutant miR-200a/b binding sites for wild-type, *Zeb1*$^{200H}$, and *Zeb1*$^{200M}$ pituitary (n = 3–4 per genotype). Each bar represents a different animal, and the total number of reads are indicated in white.

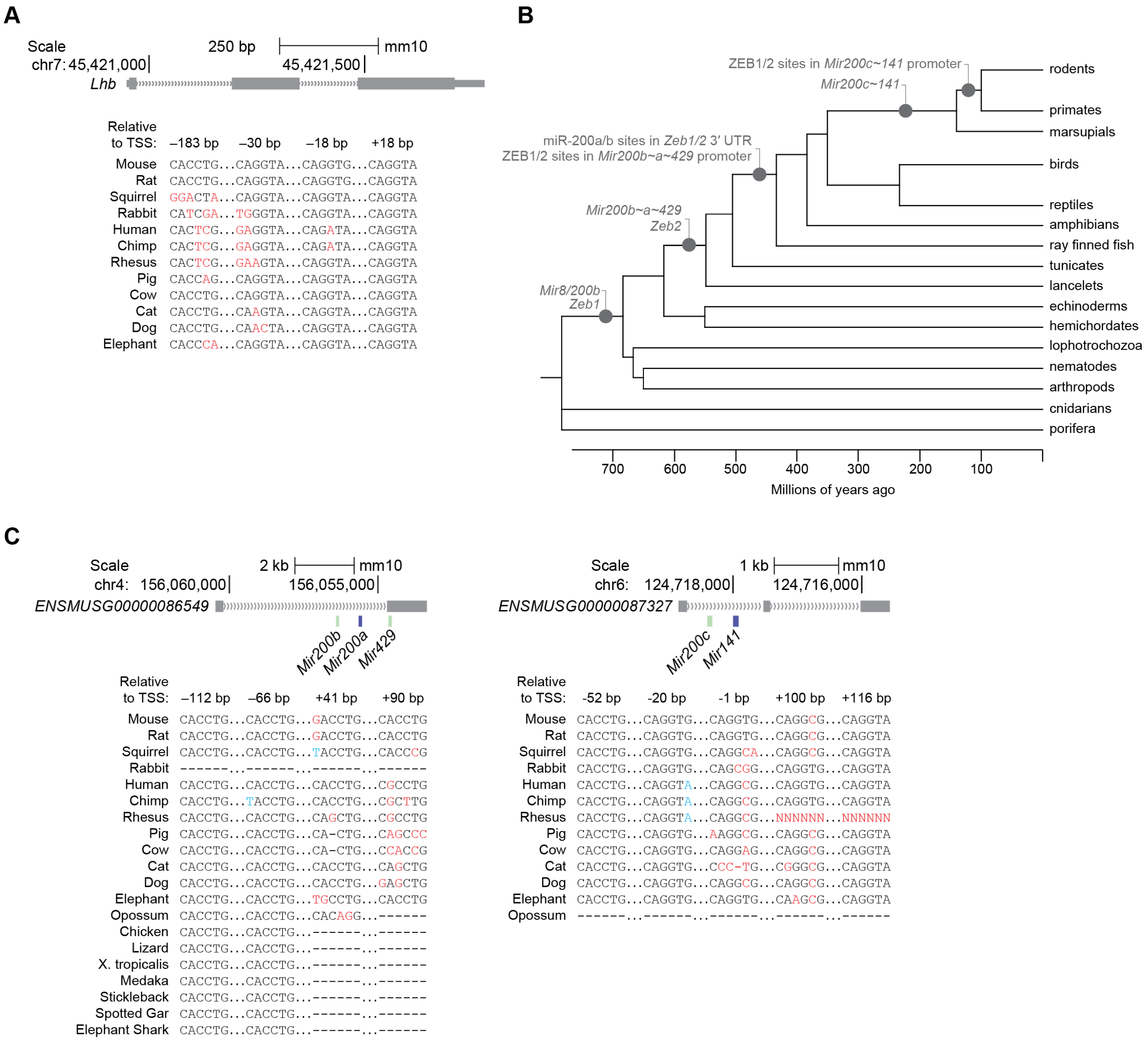

**Figure S5. Conservation of the miR-200a/b–ZEB1/2 DNFL.** (A) Organization of the murine *Lhb* locus. The *Lhb* gene model (gray boxes, exons; > > >, introns) is depicted. Diagramed below are four putative ZEB1/2 binding sites, their position relative to the mouse TSS, and Multiz alignments for each site in 12 placental mammal species (red denotes a nucleotide change that disrupts the CAGGTR consensus sequence). Not pictured are two additional human- or primate-specific CAGGTR motifs located at –604 bp and +152 bp relative to the human TSS. (B) Phylogeny of Metazoa annotated with likely introduction of genes and regulatory sequences in the miR-200a/b–ZEB1/2 DNFL. (C) Organization of the murine *Mir200b~a~429* (left) and *Mir200c~141* (right) loci. The transcript model of the presumptive murine host gene for each miRNA cluster (gray boxes, exons; > > >, introns) is depicted with the miRNA hairpins shown below (green and blue boxes). Diagramed below that are four or five putative ZEB1/2 binding sites, their position relative to the mouse TSS, and Multiz alignments for each site in 13 mammals and, for *Mir200b~a~429*, seven more deeply branching vertebrates (red denotes a nucleotide change that disrupts the CAGGTR consensus sequence, blue denotes a nucleotide change that does not disrupt the CAGGTR consensus).
